# DNA template heterogeneity and *in vitro* transcription reaction conditions impact the poly(A) tail length and heterogeneity of mRNA

**DOI:** 10.64898/2026.07.02.735822

**Authors:** Gareth R. Owen, Caroline A. Evans, Adithya Nair, Sebastian J. Ross, Mollie A. Glenister, Zoltán Kis, Mark J. Dickman

## Abstract

mRNA technology has emerged as a powerful new class of medicines. Importantly, this RNA-based approach holds promise for treatments beyond vaccines and infectious diseases, including treatments for cancer, metabolic disorders, cardiovascular conditions and autoimmune diseases. The 3’-polyadenylated (poly(A)) tail of mRNA is required for ribosome initiation, translation, and mRNA stability and is considered a critical quality attribute. In this study, novel direct mass spectrometry approaches were used for the analysis of both the DNA template and corresponding mRNA generated via *in vitro* transcription. Nucleotide resolution of the poly(A/T) sequence of the DNA template and mRNA poly(A) tail was achieved. The results show that the mRNA poly(A) tail length and heterogeneity is impacted by the heterogeneity of the DNA template, the DNA template design and RNA manufacturing conditions, including relative NTP concentrations. These results provide further important mechanistic insight into the poly(A) tail length and heterogeneity of mRNAs synthesised *in vitro*, including the identification of 3’-end additions of cytidine to mRNA poly(A) tails. The ability to rapidly assess DNA template quality, combined with monitoring mRNA poly(A) tail length and heterogeneity, is important as part of the characterisation of mRNA precision medicines and ensuring consistent quality of mRNA from manufacturing processes.

## INTRODUCTION

The development and global application of two pioneering mRNA vaccines, Comirnaty (Pfizer/BioNTech) and Spikevax (Moderna) have exemplified the power of mRNA medicine and fuelled their further development (1–3). mRNA-based approaches have potential for treatments for infectious diseases, cancer, metabolic disorders, cardiovascular conditions, and autoimmune diseases (4–7). mRNA-based medicines work by translating exogenous mRNA into the target protein, which was first demonstrated *in vivo* in 1990 by Wolff *et al*., where functional protein expression was demonstrated after direct injection of mRNA (7).

mRNA medicines contain five essential components, including: 1) 5ʹ-cap, which is required for ribosome initiation, translation, and stability (8); 2) 5ʹ-untranslated region (UTR), that helps drive high levels of translation from the correct start codon and confer mRNA stability; 3) Coding sequence, encoding for the gene of interest; 4) 3ʹ-UTR, required for translation and stability and finally; 5) 3’-polyadenylated (poly(A)) tail, required for ribosome initiation, translation, and mRNA stability (9–11). In addition, various chemical modifications (e.g. N1-methylpseudouridine) have been introduced into the mRNA to reduce immunogenicity and enhance stability and protein expression (12–14).

The 3’-poly(A) tail is typically incorporated into the initial plasmid DNA for transcription. However, mRNA can also be synthesised without a 3’-poly(A) tail by using a “tailless” pDNA template, followed by a post-transcriptional poly(A) tailing step (15, 16). In this case, poly(A) polymerase is used to add poly(A) tails of commonly 80–160 nucleotides. Poly(A) tails can be intact sequences of repeated adenosine nucleotides or segmented, i.e. ‘split’ by spacer elements, designed to mitigate technical challenges associated with plasmid DNA encoded templates for *in vitro* transcription (IVT) (17).

The length of the poly(A) tail affects the stability and translational efficiency of the mRNA. The impact of poly(A) tail length on expression and stability has been extensively studied, determining the optimum length to minimise mRNA decay caused by tail shortening and hyperadenylation (9, 18). The systematic substitution of non-A nucleotides in the tails revealed that cytidine-containing tails can substantially enhance the protein production rate and duration of synthetic mRNAs in cell lines and a mouse model (19). Chemical modification of the 3’-poly(A) tail provides a strategy to reduce or prevent their degradation (20–22). Click chemistry approaches for ‘modular’ addition of chemical modifications to the 3’-poly(A) tail, including the addition of chemically modified synthetic oligonucleotides (21) or trimers of modified poly(A) tail via a branched topology have been developed (22). Both the 5ʹ-cap and 3’-poly(A) tail affect binding to the ribosomal machinery and mRNA stability (10). Therefore, the percentage of 5’ capped mRNA (5’ capping efficiency) and the 3’-poly(A) tail length and heterogeneity are critical quality attributes (CQAs) that will impact the translational efficiency and *in vivo* stability of the mRNA (9, 18, 22). Therefore, analytical methods for the characterisation of mRNA drug substances and drug products must be developed and performed to monitor these CQAs.

### Characterisation of 3’-poly(A) tails length and heterogeneity

Assessment of the poly(A) tail involves measuring its length, which typically ranges from 80-160 nt, as well as evaluating its heterogeneity. In the context of 3’-heterogeneity, this variation can arise from transcriptional slippage, particularly in regions containing repeat mononucleotide sequences. This phenomenon can lead to the incorporation of additional nucleotides, resulting in heterogeneity (23–26). There are several high-throughput methods for detecting and profiling poly(A) tails, including TAIL-seq (27), full-length mRNA sequencing (FLAM-seq (28)) and poly(A) inclusive RNA isoform sequencing (PAIso-seq (29)). A major caveat is that they require an amplification step, which leads to variations in sequencing. Direct RNA sequencing (DRS) from Oxford Nanopore Technologies eliminates potential sources of amplification bias and has been widely employed for poly(A) profiling. However, the homopolymer stretches in DRS cannot be precisely base-called due to the uniformity of the raw signal between adjacent nucleotides (30–33). Therefore, using such approaches typically enables only an average poly(A) tail length to be estimated. More recently, direct nanopore sequencing of synthetic mRNA molecules in conjunction with computational methods was used to detect mis-inserted non-adenosine residues in mRNA poly(A) tails (34).

Mass spectrometry methods for the characterisation of the mRNA poly(A) tail were first developed by Beverly *et al*. to characterise IVT-synthesised mRNA with varying length poly(A) tails (35). RNase T1 digests in conjunction with oligo dT purification of the poly(A) tail fragment prior to LC-MS analysis enabled nucleotide resolution of the mRNA poly(A) tail, demonstrating lengths greater than expected from the corresponding DNA template (35). More recently, the advantages of mass spectrometry analysis of the DNA template using PCR prior to LC-MS analysis further highlighted the advantages of LC-MS-based approaches for the analysis of polypurine/pyrimidine sequences in DNA templates compared to Sanger sequencing (36).

In this study, direct LC-MS approaches were used for the analysis of both the DNA template and the corresponding mRNA generated via IVT. Nucleotide resolution of the poly(A/T) sequence of the DNA template and mRNA poly(A) tail shows that the mRNA poly(A) tail length and heterogeneity are impacted by the heterogeneity of the DNA template, the DNA template design and manufacturing conditions, including relative NTP concentrations.

## RESULTS

### DNA template heterogeneity affects mRNA poly(A) tail length and heterogeneity

The effect of the DNA template on the length and heterogeneity of the poly(A) tail of in vitro transcribed mRNA was studied through the analysis of multiple DNA templates and their resulting mRNA transcripts. A wide range of different DNA templates with a poly(A) tail encoded on the DNA template were obtained from both commercial suppliers and in-house plasmid cloning (see Table 1). Verification of the DNA template sequence and characterisation of the poly(A/T) region was performed using both Sanger sequencing and a novel LC-MS-based approach to directly study the poly(A/T) sequence of the DNA template in conjunction with restriction digestion to determine the relative abundance and distribution of individual polynucleotide species. mRNA transcribed from these DNA templates was also analysed by LC-MS, following RNase T1 digestion, to study the resulting mRNA poly(A) tail length and heterogeneity.

**Table 1.** Summary of DNA templates used in this study. DNA templates and corresponding mRNAs are shown with the predicted mRNA size and poly(A) tail length as determined by Sanger sequencing during template production. ^*^ Expected mRNA size with expected poly(A) tail, † Predicted poly(A) tail length (Sanger sequencing).

| DNA Template<br>Number | Template Type | mRNA<br>Sequence | mRNA Size<br>(nt)* | Poly(A) Tail<br>Length (nt)† |
| --- | --- | --- | --- | --- |
| 1 | Linearised Plasmid | NLuc | 871 | 120 |
| 2 | Linearised Plasmid | eGFP | 930 | 49 |
| 3 | Linearised Plasmid | CSP | 4286 | Segmented 30/70 |
| 4 | Linearised Plasmid | eGFP | 995 | 105 |
| 5 | PCR Product | eGFP | 930 | 49 |
| 6 | Enzymatically<br>synthesised DNA | eGFP | 1168 | 94 |

A linearised plasmid DNA template encoding a NanoLuc Luciferase mRNA was purchased from a commercial supplier with an expected 120 nt poly(A) tail. Sanger sequencing of the plasmid DNA template is shown in Figure 1a, which reported a DNA poly(A) sequence of approximately 115 nts before confident base assignment was not possible due to heterogeneous sequencing reads. Moreover, the sequencing of the (antisense) strand reported a different length of the complementary poly(T) sequence (118 nts) (See Supplementary Figure S1). These results highlight the caveats of Sanger sequencing approaches, which are unable to accurately determine the length of polypyrimidine/purine sequences and analyse potentially heterogeneous DNA sequences (35, 36). Sanger sequencing data after these homopolymeric sequences were characterised by low-quality and low-confidence reads. Therefore, to further characterise the plasmid DNA template, the plasmid DNA was restriction digested to generate a DNA fragment that encompasses the poly(A/T) region, prior to direct analysis using LC-MS. The digested plasmid DNA was separated using ion-pair reversed-phase HPLC (IP-RP HPLC) online to an Orbitrap Exploris mass spectrometer. During the chromatographic separation, the poly(A) and poly(T) strands of the DNA duplex separate from each other and are resolved from the other template digest fragments (See Supplementary Figure S2), mitigating the need for further purification of the fragments of interest. Intact mass deconvolution was used across the retention times of the chromatographic peaks corresponding to the poly(A) and poly(T) containing strands to determine the accurate mass of the DNA species present. Following intact mass deconvolution, the corresponding length of the poly(A) and poly(T) ssDNA was determined with nucleotide resolution (see Figure 1b). The results show that the most abundant length species for the sense and antisense strands of the poly(A/T) fragment was 118 nts. Moreover, the mass spectrometry analysis revealed a heterogeneous population in the DNA template, ranging from 95-121 nts, with three major populations observed (see Figure 1b). These results highlight the advantages of characterising the poly(A/T) region of DNA templates designed for the manufacturing of mRNA using LC-MS in comparison to Sanger sequencing. The LC-MS analysis reveals significant heterogeneity of the poly(A/T) region of this commercially obtained DNA template. Following characterisation of the DNA template, the mRNA was transcribed using IVT prior to RNase T1 digestion and direct LC-MS analysis of the resulting poly(A) tail. Following intact mass deconvolution, the corresponding length of the poly(A) tail was determined with nucleotide resolution (see Figure 1c). The results show that mRNA with a 120 nt poly(A) tail as the most abundant species. Moreover, the data demonstrated a multimodal distribution, which mirrors the corresponding DNA template (Figure 1c). This suggests that although there is some transcriptional slippage of the T7 RNA polymerase that results in the production of longer poly(A) tail lengths from the corresponding DNA template, the majority of the heterogeneity in the mRNA can be attributed to heterogeneity present in this particular heterogeneous plasmid DNA template.

**Figure 1.**
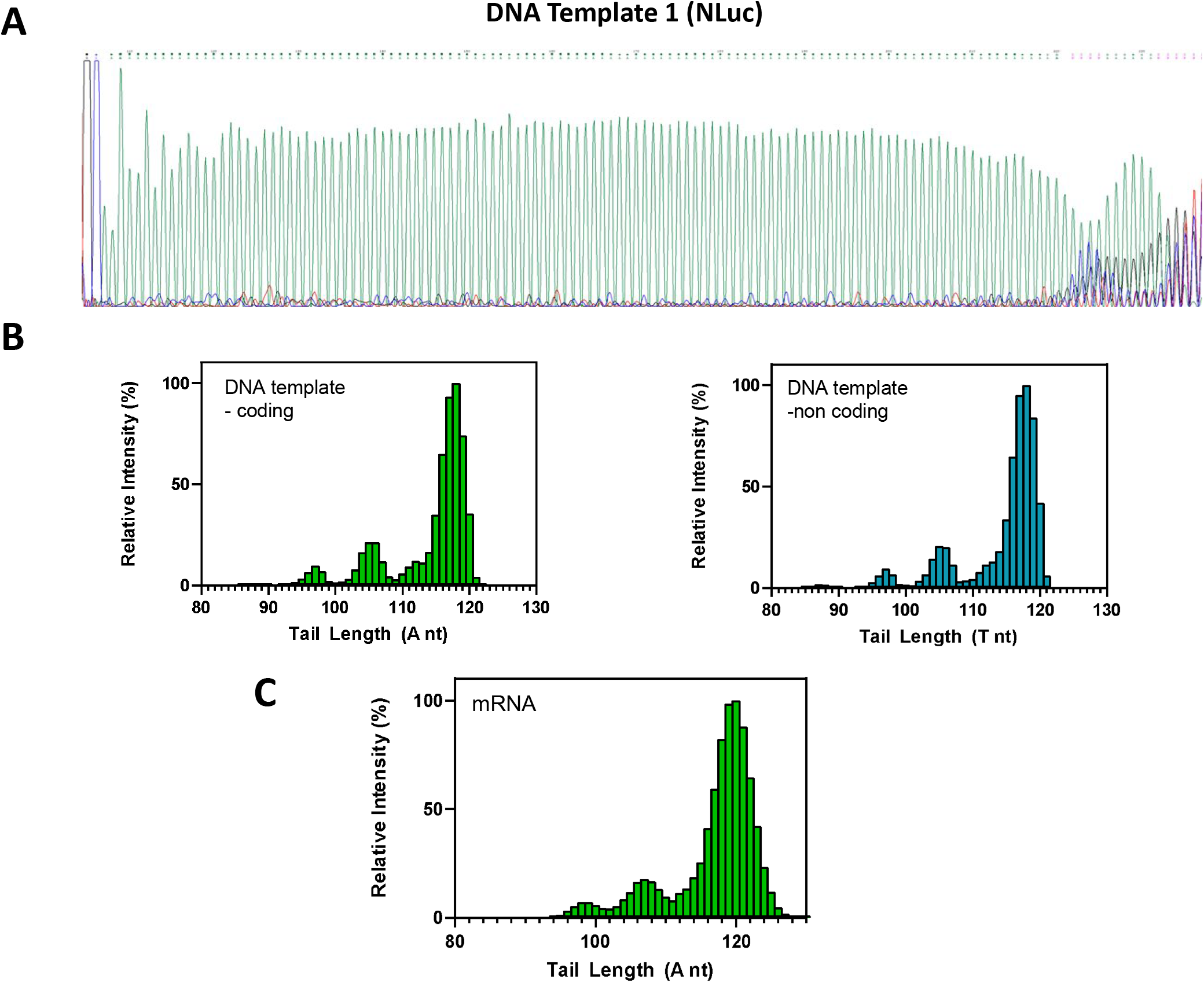
Characterisation of DNA and mRNA poly(A) tail length and heterogeneity of template 1 encoding an NLuc mRNA. (A) A representative Sanger sequencing chromatogram of the poly(A) encoding region of the DNA template used for *in vitro* transcription. (B) LC-MS analysis and identification of the coding/poly(A) and non-coding/poly(T) strands from restriction digestion of the DNA template. (C) mRNA poly(A) tail species identified by LC-MS from RNase T1 digestion of *in vitro* transcribed mRNA.

Further analysis was performed using an alternative commercially obtained DNA template encoding the COVID Spike protein mRNA (CSP mRNA). The DNA template was synthesised with a split poly(A/T) region with a proposed 70/30 nt split poly(A) tail with an intervening 10 nt linker . Split poly(A) tails have been utilised in an approach to enhance the stability of the poly(A/T) region and minimise recombination of plasmid DNA (17, 39). Sanger sequencing of the DNA template covering the poly(A/T) region is shown in Figure 2a. The results show the expected poly(A) length of 30 nts followed by the 10 nt linker and the final poly(A) sequence of 69 nts. Similar to previous Sanger sequencing, heterogeneity was observed in the sequencing data downstream of the poly(A) tract with the 70 nt position a mixed base called limiting accurate characterisation of the exact size and heterogeneity present in the DNA template. In addition, LC-MS analysis of the poly(A/T) region was performed using restriction digestion of the DNA template as previously described (see Figure 2b). The results confirm the poly(A/T) sequence lengths as 30 nts and 70 nts as the most abundant species for the two ssDNA strands, with no significant heterogeneity of the homopolymeric A and T regions (<2% relative intensity). mRNA was subsequently transcribed from the corresponding DNA template prior to intact mass analysis of the poly(A) tail as previously described (see Figure 2c). The results show a narrow distribution of the length of the poly(A) tail, ranging from 67-77 nts, with the most abundant species having a 71 nt poly(A) tail. These results demonstrate, in this case, that the heterogeneity of the poly(A) tail arises predominantly through transcriptional slippage and the addition of adenosine nucleotides beyond the expected 70 nt from the corresponding DNA template. Only minor species are observed <70 nts, which would likely correspond to transcriptional slippage of the T7 polymerase on the poly(T) sequence of the DNA template, resulting in the deletion of nucleotides from the poly(A) tail (Figure 2c).

**Figure 2.**
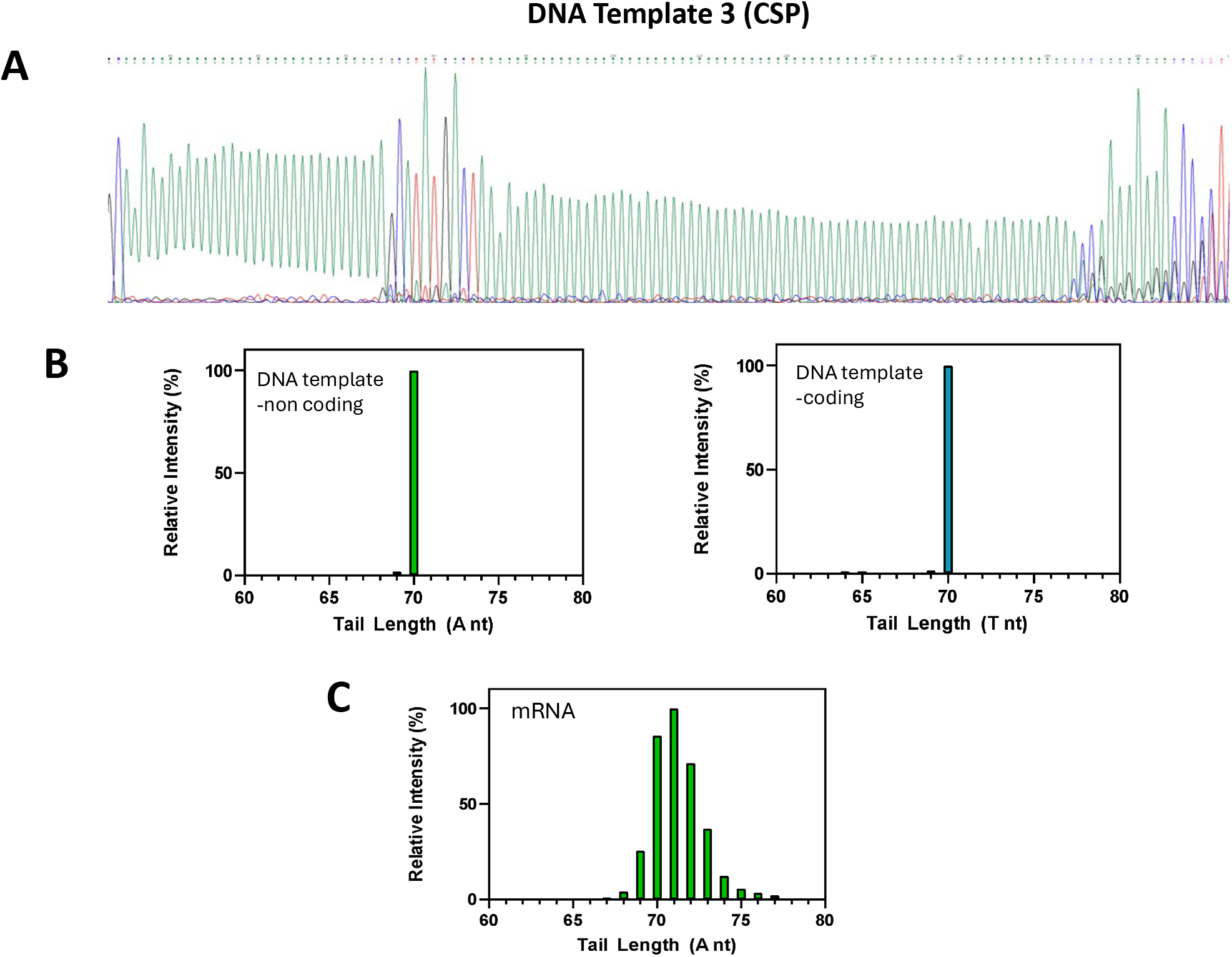
Characterisation of DNA and mRNA poly(A) tail length and heterogeneity of template 3 encoding a COVID Spike Protein mRNA. (A) A representative Sanger sequencing chromatogram of the poly(A) encoding region of the DNA template used for *in vitro* transcription. (B) LC-MS analysis and identification of the coding/poly(A) and non-coding/poly(T) strands from restriction digestion of the DNA template corresponding to the 70 nt region(C) mRNA poly(A) tail species identified by LC-MS from RNase T1 digestion of *in vitro* transcribed mRNA corresponding to the 70 nt region.

In addition to the characterisation and analysis the poly(A) tail from IVT-generated mRNA from commercially obtained plasmid DNA templates, we generated plasmid DNA constructs using a pre-linearised vector and eGFP coding sequence. Validation and characterisation of the recombinant plasmid DNA was performed using Sanger sequencing (see Figure 3). The results show a poly(A/T) sequence length of 104 nts followed by heterogeneous sequence data at the following positions (Figure 3a). Restriction digestion and LC-MS analysis identified the length of poly(A/T) sequence as 105 nts, with only minor sequence lengths <2% relative intensity detected (see Figure 3b). mRNA transcribed from the linearised DNA template resulted in the synthesis of mRNA with a poly(A) tail length of 106 nt as the most abundant species, with a narrow distribution of shorter and longer tails (see Figure 3c). These results also demonstrate that the heterogeneity of the poly(A) tail arises predominantly through transcriptional slippage and the addition of adenosine nucleotides beyond the expected size from the corresponding DNA template with only minor shorter poly(A) species via transcriptional slippage of the T7 polymerase on the DNA template, leading to the deletion of nucleotides.

**Figure 3.**
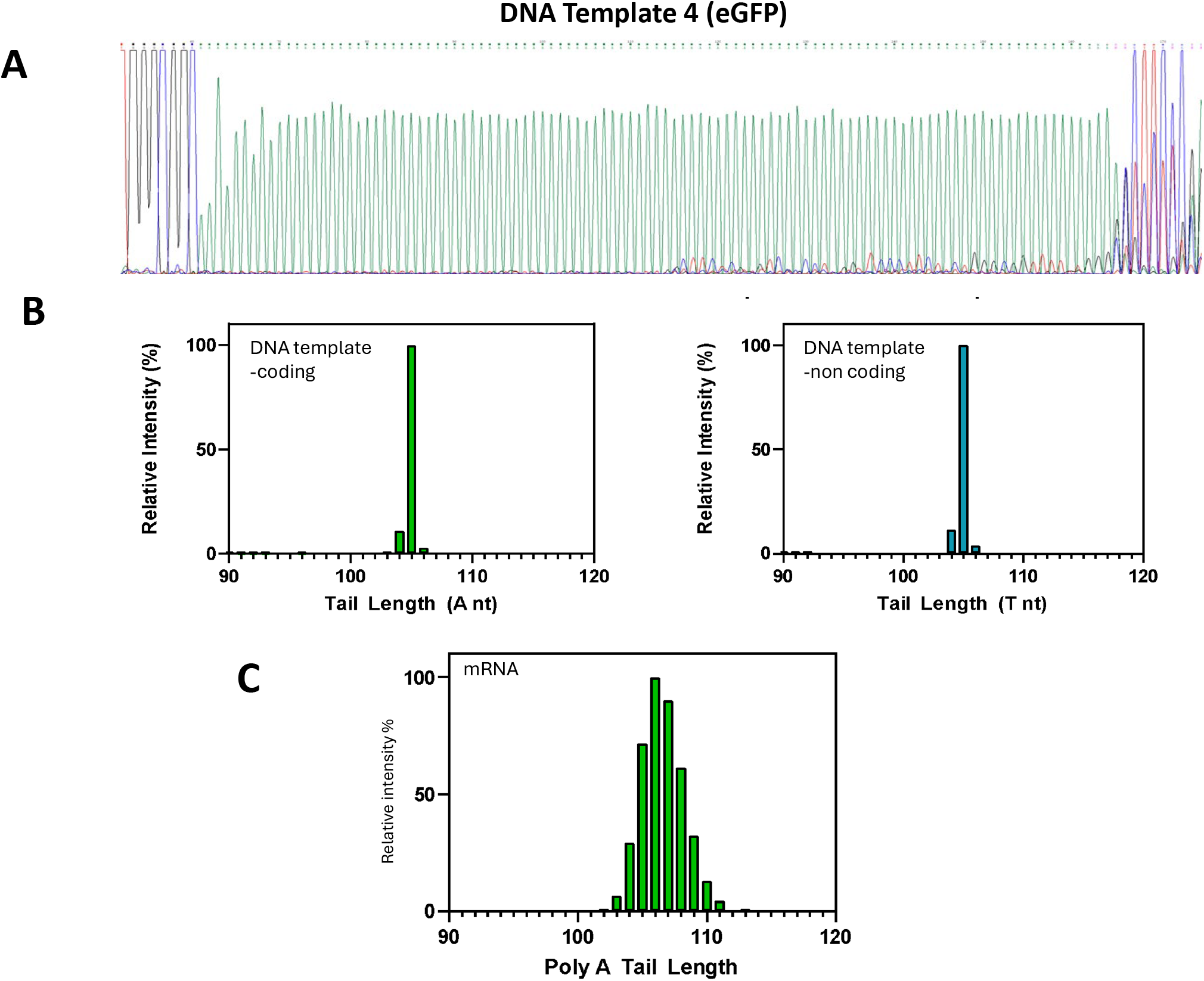
Characterisation of DNA and mRNA poly(A) tail length and heterogeneity of template 4 encoding an eGFP mRNA. (A) A representative Sanger sequencing chromatogram of the poly(A) encoding region of the DNA template used for *in vitro* transcription. (B) LC-MS analysis and identification of the coding/poly(A) and non-coding/poly(T) strands from restriction digestion of the DNA template. (C) mRNA poly(A) tail species identified by LC-MS from RNase T1 digestion of *in vitro* transcribed mRNA.

### DNA linearisation strategies affect poly(A) tail length and heterogeneity of mRNA

Previous analysis demonstrated the effects of heterogeneity of the poly(A/T) sequence in the DNA template on the mRNA poly(A) tail and the ability of LC-MS methods to accurately measure DNA template heterogeneity. In the previous examples, all linearised DNA templates had additional nucleotides downstream of the poly(T) sequence. Previous results have indicated that the addition of nucleotides downstream of the poly(T) sequence can influence poly(A) tails in IVT reactions (35). Therefore, further studies were performed using LC-MS analysis of mRNA poly(A) tails generated from linearised DNA templates in the absence of additional nucleotides downstream of the poly(T) sequence at the 5’-end of the DNA template.

A DNA template encoding eGFP mRNA with an expected 49 nt poly(A) tail was analysed using both Sanger sequencing and LC-MS (see Figure 4a/b). The Sanger sequencing showed the expected 49 nt poly(A) length with only limited signal heterogeneity downstream of the poly(A) sequence (Figure 4a). LC-MS analysis of the poly(A/T) identified a sequence of 49 nts as the most abundant length, with effectively no heterogeneity in the DNA template (>2% relative intensity) (see Figure 4b). Prior to IVT, the DNA template was either linearised with either SfiI which cuts after the poly(T) sequence and generates mRNA with additional nucleotides downstream of the poly(A) tail or BspQI which cuts within the poly(T) sequence and therefore generates mRNA with no additional nucleotides downstream of the poly(A) tail. The corresponding LC-MS analysis of the mRNA poly(A) tails is shown in Figures 4c/d.

**Figure 4.**
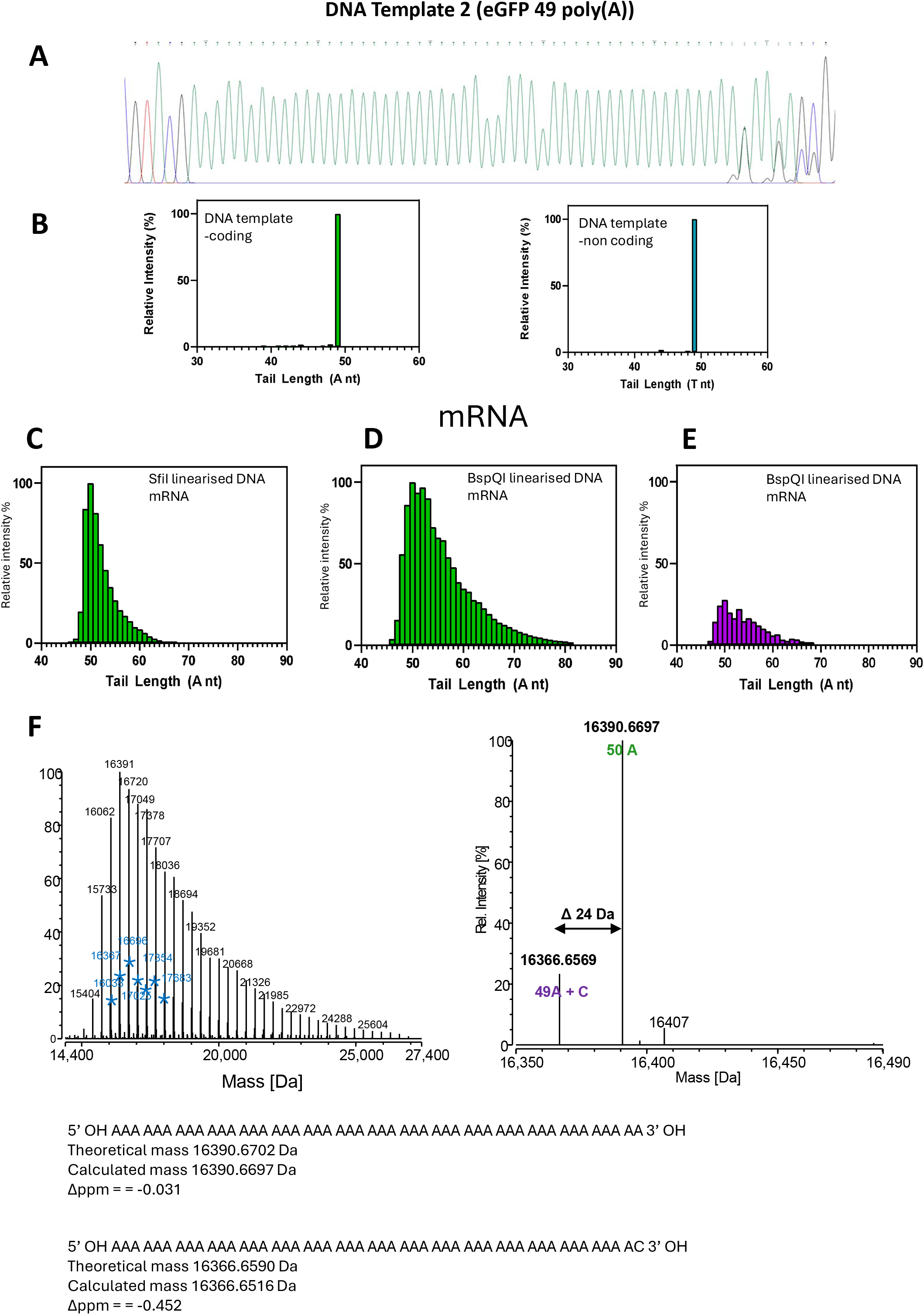
Characterisation of DNA and mRNA poly(A) tail length and heterogeneity of template 2 encoding an eGFP mRNA. (A) A representative Sanger sequencing chromatogram of the poly(A) encoding region of the DNA template used for *in vitro* transcription. (B) LC-MS analysis and identification of the poly(A) and poly(T) strands from restriction digestion of the DNA template. mRNA poly(A) tail species identified by LC-MS from RNase T1 digestion of *in vitro* transcribed mRNA from template linearised with either SfiI (C) or BspQI (D), where an additional cytidine-containing population is observed (E). The relative abundance of the poly(A) tail population is shown in green and poly(A) tail +C population is shown in magenta (F) Deconvoluted spectra from the high-resolution accurate mass analysis demonstrating the identification of the two tail populations.

The results show clear differences in the length and heterogeneity of the mRNA poly(A) tail from the two DNA templates. The results show that a wider distribution of longer poly(A) tail lengths was observed from DNA templates with no additional nucleotides present after the poly(T) sequence at the 5’-end of the DNA template. These results indicate an increase in transcriptional slippage and non-templated addition of A nucleotides into the mRNA poly(A) tail, consistent with previous observations of T7 RNA polymerase (23–25, 35). Furthermore, the LC-MS analysis of mRNA generated from linearised DNA templates that have no additional nucleotides downstream of the DNA poly(T) sequence enables further analysis of any potential heterogeneity that may arise from 3’-end additions of the mRNA when using RNase T1 digests prior to LC-MS analysis. Further analysis of the LC-MS data revealed two distinct populations based on the high-resolution accurate mass analysis (see Figure 4d/e). Based on the mRNA sequence, the experimental masses correspond to the RNase T1 fragment, including the poly(A) tail with an additional cytidine present on each of the poly(A) mRNA tails (see Figure 4f). The relative intensity of this population was lower than the poly(A) tail with no 3’-end additions (see Figure 4d/e).

Further analysis of the mRNA poly(A) tail generated from a similar DNA template, which was also linearised within the poly(T) sequence, is shown in Supplementary Figure S3. In this case, a commercial enzymatically synthesised DNA template was obtained. The Sanger sequencing and LC-MS analysis of the poly(A/T) sequence show significant heterogeneity in this DNA template. The resulting mRNA was found to have a 95 nt poly(A) tail as the most abundant species with significant heterogeneity reflecting the heterogeneity in the DNA template and additional transcriptional slippage resulting in longer length poly(A) tails. Consistent with the previous data, the LC-MS analysis also identified a secondary lower abundance population of poly(A) tails consistent with the 3’-end addition of cytidine.

### PCR amplification of the DNA template results in increased heterogeneity in the poly(A/T) sequence

In addition to the analysis of the DNA template poly(A/T) sequence using direct restriction enzyme digestion and LC-MS analysis, the DNA template corresponding to eGFP was amplified using PCR prior to Sanger sequencing and restriction digestion/LC-MS analysis (see Figure 5). The results show that following amplification from the original DNA template, an increase in heterogeneity of the poly(A/T) sequence was observed (see Figure 4 and Figure 5). These results demonstrate that during amplification, additional heterogeneity is introduced, potentially arising through slippage of the DNA polymerase in the poly(A/T) sequence (40–43). These results further highlight the advantages of directly analysing the DNA template via restriction digestion and LC-MS to accurately characterise the poly(A/T) sequence, compared to PCR amplification approaches, which result in increased heterogeneity of the DNA template. The corresponding mRNA from the PCR amplified DNA template was generated using IVT and digested using RNase T1 prior to LC-MS analysis of the corresponding mRNA poly(A) tail (see Figure 5c).

**Figure 5.**
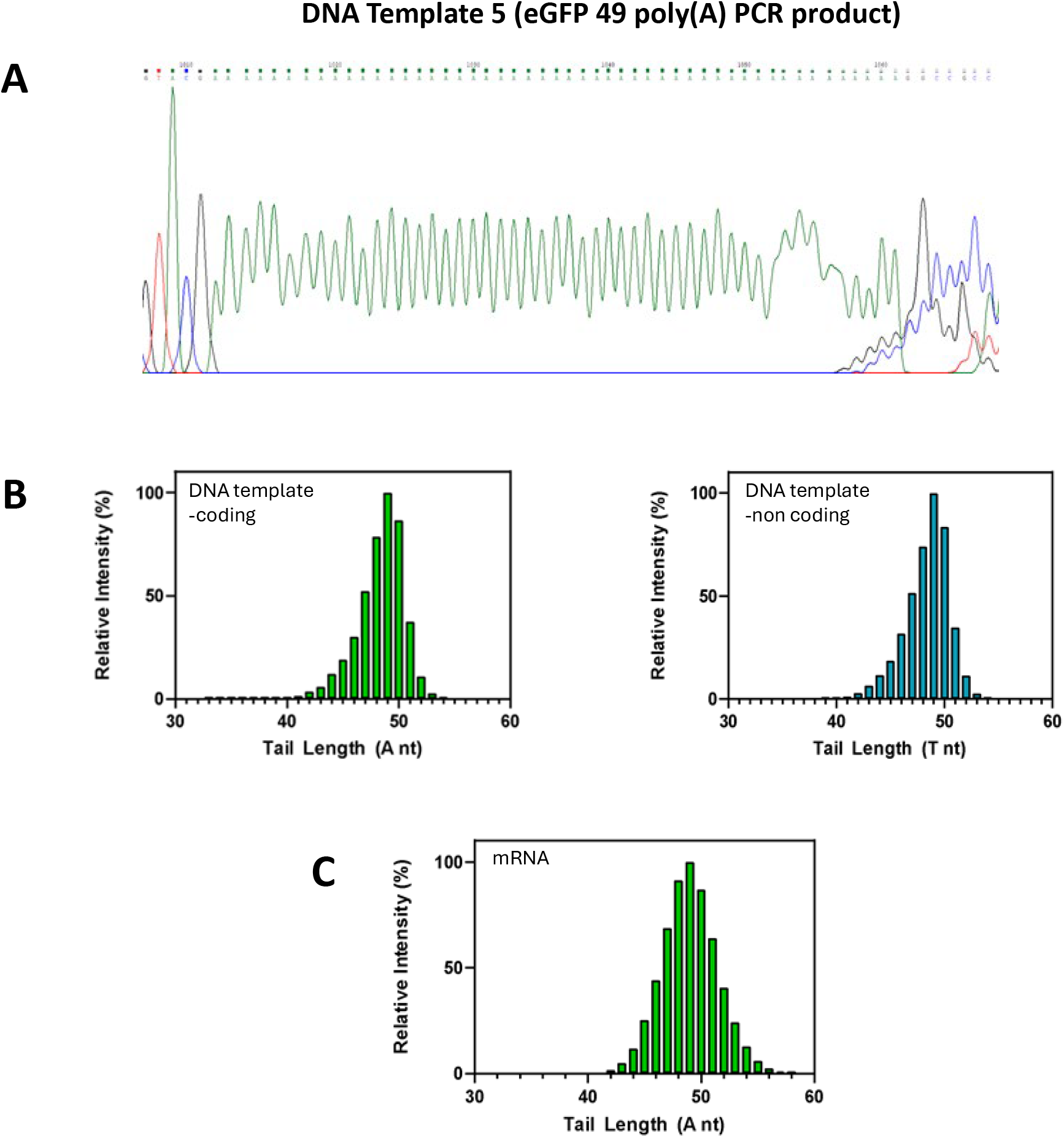
Characterisation of DNA and mRNA poly(A) tail length and heterogeneity of template 6 encoding eGFP mRNA. (A) A representative Sanger sequencing chromatogram of the poly(A) encoding region of the DNA template used for *in vitro* transcription. (B) LC-MS analysis and identification of tail species from the coding/poly(A) and non-coding/poly(T) strands from restriction digestion of the DNA template. (C) mRNA poly(A) tail species identified by LC-MS from RNase T1 digestion of *in vitro* transcribed material.

### IVT manufacturing conditions impact mRNA poly(A) tail length and heterogeneity

#### Studying the effect of ATP concentration

To study the role of ATP concentration on the mRNA poly(A) tail length and heterogeneity, IVT reactions were performed from a range of different linearised DNA templates previously characterised using LC-MS. Linearised DNA templates were selected with and without additional nucleotides at the 5’-end of the poly(T) sequence on the DNA template. IVT reactions were performed varying the ATP concentration from 0.5 mM to 15 mM, whilst the remaining NTPs were kept constant at 10 mM. Following IVT, the mRNA was digested with RNase T1 prior to LC-UV and LC-MS analysis of the corresponding mRNA poly(A) tail. The results from the initial LC-UV analysis of the mRNA poly(A) tail from DNA template 2 (eGFP) is shown in Figure 6. Due to the smaller poly(A) tail of this mRNA, the LC-UV analysis enables resolution of the different-sized poly(A) tails. The smaller poly(A) tails elute prior to the longer poly(A) tail species and therefore provide a rapid comparative analysis across the different IVT reactions. The LC-UV results show clear differences across the range of ATP concentrations used (see Figure 6a) and show a trend of increasing abundance of longer and more heterogeneous populations of poly(A) tail at higher ATP concentrations. Further comprehensive analysis of the poly(A) tail populations was performed using LC-MS (see Figure 6b). The results are consistent with the LC-UV analysis and show increasing poly(A) tail length and heterogeneity as the concentration of ATP is increased in the IVT reaction. These results demonstrate that increased transcriptional slippage is observed at high ATP concentrations, leading to longer and more heterogeneous mRNA poly(A) tails from linearised DNA templates with no additional nucleotides downstream of the poly(T) sequence.

**Figure 6.**
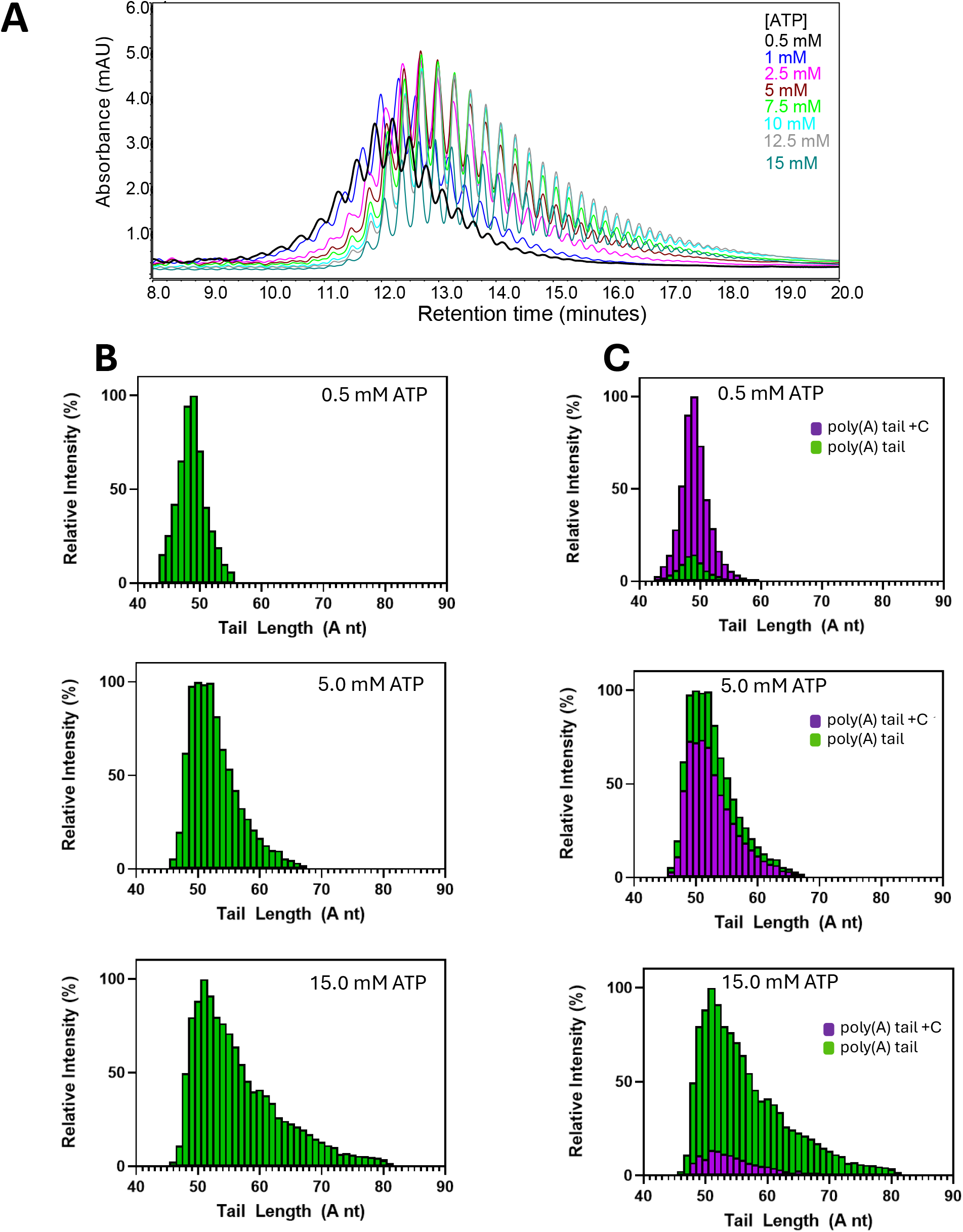
ATP concentration effects on the poly(A) tail distribution of mRNA transcribed from DNA template 2 encoding an eGFP mRNA. (A) LC-UV chromatogram of the 3’-poly(A) tail fragment generated from RNase T1 digestion of *in vitro* transcribed mRNA using varying concentrations of ATP. (B) mRNA poly(A) tail species identified by LC-MS from RNase T1 digestion of *in vitro* transcribed material using 0.5 mM, 5 mM, or 15 mM ATP. (C) The two poly(A) tail populations: ‘poly(A) +C’ (purple) or ‘poly(A)’ (green).

Furthermore, consistent with previous LC-MS analysis of mRNA transcribed from this DNA template, an additional species proposed to result from 3’-end addition of cytidine was also observed (see Figure 6c and Supplementary Figure S4). Interestingly, the results show that at the lowest concentration of ATP (0.5 mM), the relative population of mRNA tails +C was the most abundant, suggesting that the 3’-end addition of cytosine is favoured when ATP is limiting. At 5.0 mM ATP, the population of poly(A) with and without the 3’-end addition is approximately equal. At high concentrations of ATP, the population of mRNA tails without the 3’-end additions is the most abundant; however, under these conditions, increased transcriptional slippage and a longer poly(A) tail distribution are observed. Further analysis, performed on RNA generated from the enzymatically synthesised DNA template, is shown in Supplementary Figure S5.

The effect of ATP concentration on mRNA poly(A) tail length and heterogeneity was also studied for alternative linearised DNA templates (1, 3) containing additional nucleotides downstream of the poly(T) sequence (see Figure 7a/b). The results are consistent with previous data, where reduced transcriptional slippage is observed compared to the previous DNA template. However, in contrast to previous data, no increase in transcriptional slippage or mRNA poly(A) tail length is observed at high ATP concentrations. It is interesting to note that at the lowest ATP concentration used, an increase in relative abundance of shorter length mRNA poly(A) tails is observed, demonstrating a potential increase in transcriptional slippage resulting in deletion of nucleotides at low ATP concentrations. In addition, as expected, lower yields of mRNA were obtained at low ATP concentrations.

**Figure 7.**
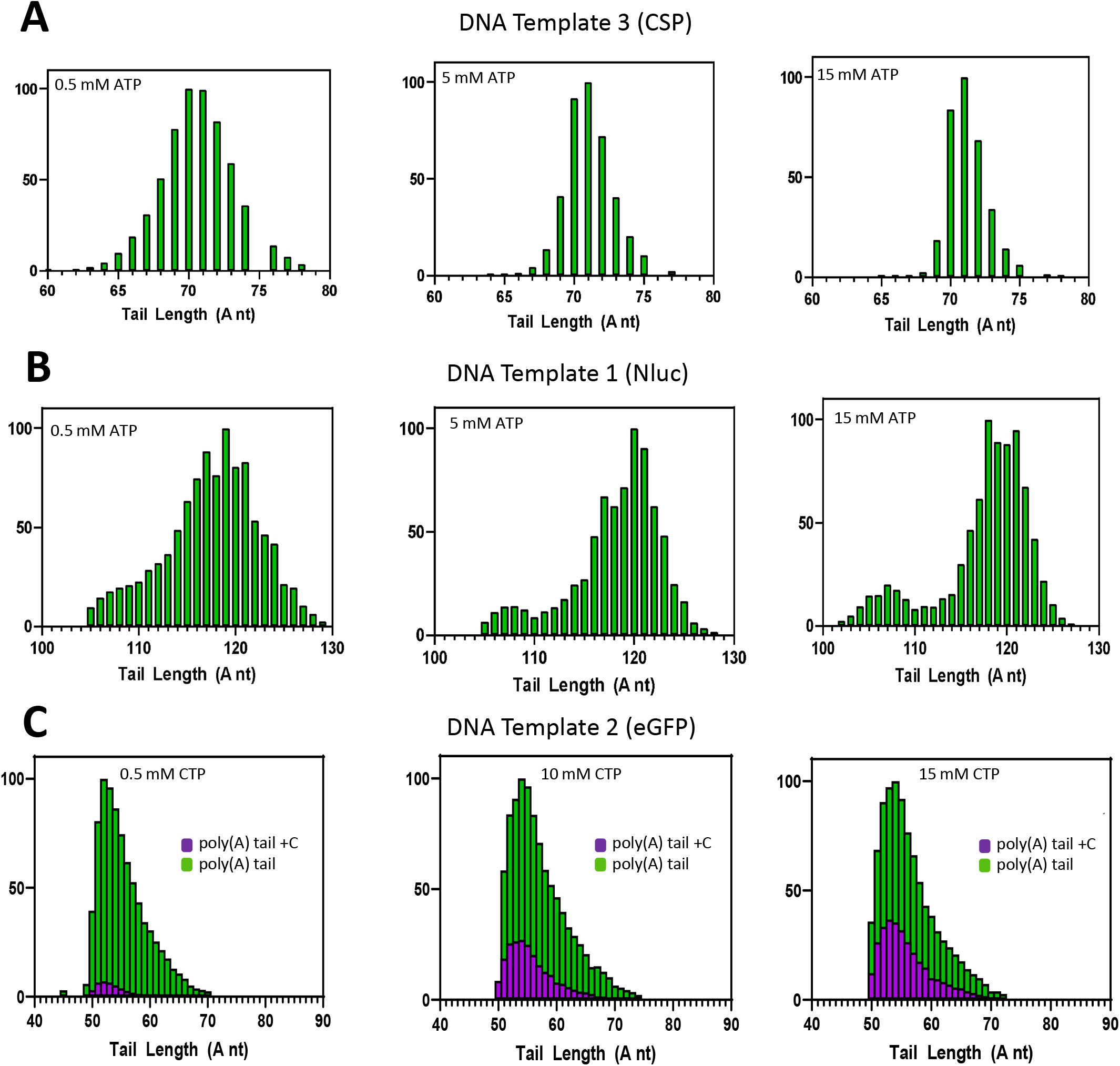
NTP concentration effects on poly(A) tail distribution. mRNA poly(A) tail species identified by LC-MS from RNase T1 digestion of *in vitro* transcribed material using 0.5 mM, 5 mM, or 15 mM ATP. A) DNA template 3 encoding a CSP mRNA (B) DNA template 1 encoding an NLuc mRNA. C) Poly (A) tail species identified by LC-MS from RNase T1 digestion of *in vitro* transcribed material using 0.5 mM, 10 mM, or 15 mM CTP from DNA template 2 encoding an eGFP mRNA.

#### Studying the effect of CTP concentration

Previous results had shown the presence of 3’-end additions of cytidine on the mRNA poly(A) tails. Therefore, to further validate these findings, the effect of CTP concentration on mRNA poly(A) tail heterogeneity was further investigated. IVT reactions were performed using DNA template 2 (eGFP) linearised with BspQI at varying concentrations of CTP, whilst remaining NTPs were constant (see Figure 7c). At all CTP concentrations studied, the most abundant population was the mRNA poly(A) tail without 3’-end additions. Furthermore, as expected, no change in the typical poly(A) tail length and heterogeneity was observed with changing CTP concentrations. However, the relative abundance of the poly(A) tail +C increased as CTP concentration increased. The results show an increase from ∼10% relative intensity at 0.5 mM CTP to ∼30% at 10 mM and ∼40% at the highest concentration of 15 mM. Therefore, these results further support the 3’-end addition of cytidine, which, as expected, is increased in abundance at high CTP concentrations in the IVT reaction. Furthermore, the effect of varying UTP concentration was also studied in conjunction with low ATP concentrations (see Supplementary Figure S6). As expected, the mRNA poly(A) tails +C were not affected by UTP concentration.

#### IVT reaction time

Further studies were performed to investigate the change in mRNA poly(A) tail length and heterogeneity over time during the IVT reaction. mRNA was produced from three different DNA templates (1-3), with mRNA taken at different time points prior to RNase T1 digestion and LC UV/MS analysis (see Figure 8). The LC-UV analysis of the poly(A) tail from DNA template 2 (eGFP) demonstrates only a small change towards longer length tails with time (Figure 8a). LC-MS analysis revealed that there was no significant change in the length and distribution of the mRNA poly(A) tail with increased reaction time. The most abundant poly(A) tail remains at 52 nts at 15, 30, and 120 minutes, respectively (Figure 8b). A small increase in the relative intensity of longer poly(A) tails at the extreme of the distribution is observed with increased IVT reaction time. In addition, a decrease in the relative abundance of the proposed 3’-end additions of cytidine to the poly(A) tail was also observed, from 34% relative intensity at 15 minutes to 24% at 120 minutes. The overall reduction in relative abundance of the mRNA 3’-end heterogeneity is proposed to be likely due to the decrease in CTP concentration in the IVT towards the end of the reaction. The impact of reaction time was further investigated with mRNA transcribed from linearised DNA templates 1 and 2, which include additional nucleotides downstream of the poly(A/T) region (see Figure 8c/d). No significant change in the relative abundance of the most abundant mRNA poly(A) tail or the overall distribution of tail lengths was observed. Overall, the data show that the IVT reaction time has a limited effect on the mRNA poly(A) tail length and heterogeneity.

**Figure 8.**
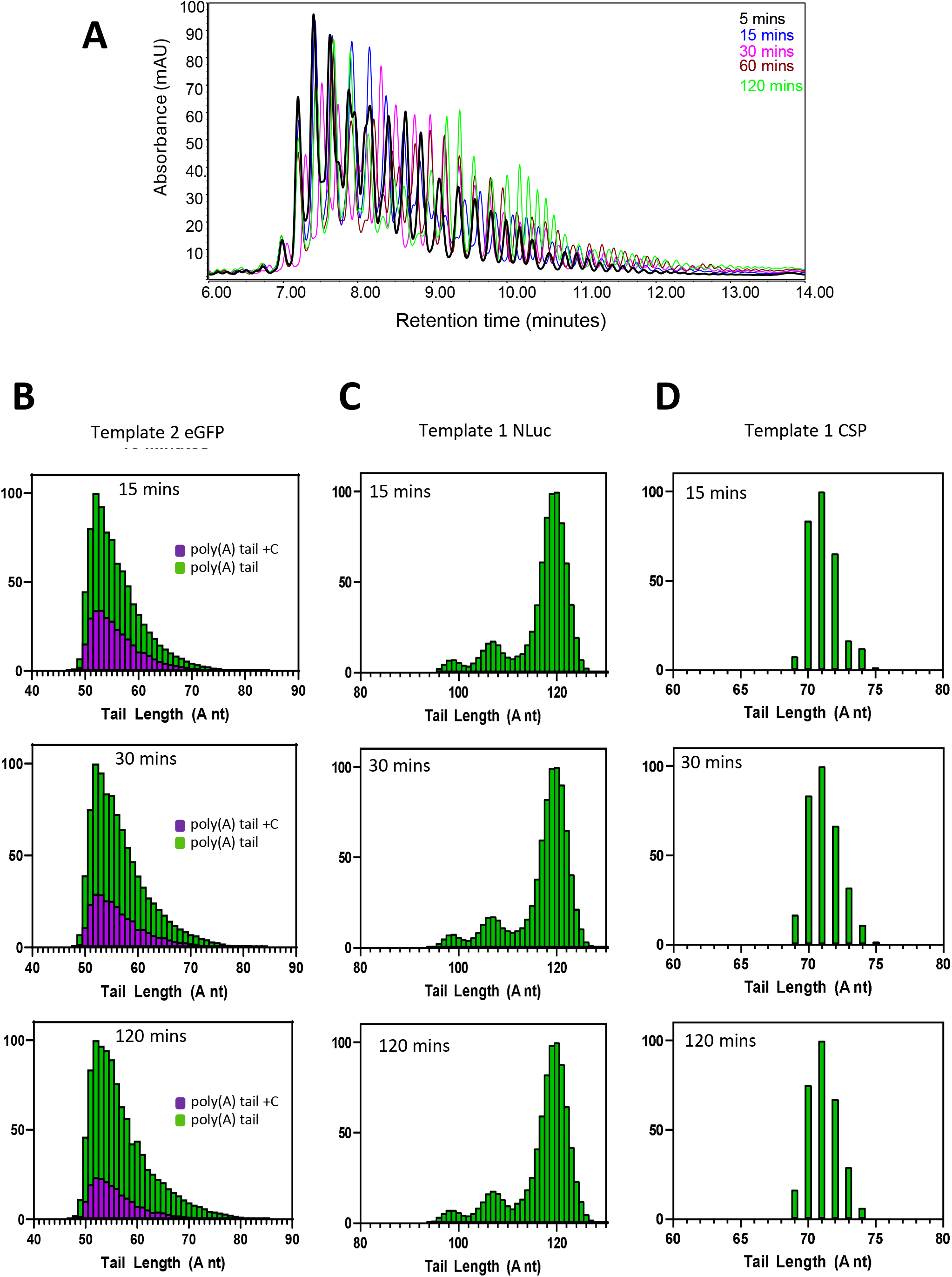
Analysis of IVT reaction time on poly(A) tail length and heterogeneity. (A) LC-UV chromatogram of the 3’-poly(A) tail from *in vitro* transcribed mRNA generated from DNA template 2 encoding an eGFP mRNA. mRNA samples were taken from a range of reaction times; 15 minute, 30 minute, and 120 minute reactions. Corresponding LC-MS analysis and identification of the 3’-poly(A) tails are shown in (B-D) for the different DNA templates. The poly(A) and poly(A)+C populations observed in mRNA transcribed from DNA template 2 are shown in green and purple respectively.

## DISCUSSION

In this study, we have developed a novel LC-MS method to directly characterise DNA plasmid templates. Restriction digestion and LC-MS analysis were used to characterise the length and heterogeneity of poly(A/T) sequences of DNA templates and therefore rapidly assess DNA template quality. The LC-MS method enables nucleotide resolution of the poly(A/T) region, demonstrating significant advantages over Sanger sequencing and NGS approaches for the characterisation of poly(A/T) regions in DNA templates used for the manufacturing of mRNA medicines. Moreover, the analysis of a wide range of commercial plasmid DNA templates and DNA templates prepared in-house demonstrates significant variation in DNA template quality with respect to the heterogeneity of the poly(A/T) sequence. Therefore, it is proposed that the direct LC-MS approaches should be used for the characterisation and quality control analysis of DNA templates used in the manufacturing of mRNA medicines, in addition to traditional Sanger sequencing and low-resolution restriction digestion and gel electrophoresis methods currently utilised.

Further studies were performed using IVT from a wide range of linearised plasmid DNA templates previously characterised using LC-MS to generate the corresponding mRNA. Following mRNA synthesis and purification, RNase T1 digests were used to generate the poly(A) tail prior to high-resolution mass spectrometry analysis. The results demonstrate that for low-quality DNA templates that exhibit heterogeneity in the poly(A/T) sequence, the corresponding mRNA poly(A) tail mirrors the heterogeneity of the DNA template. For high-quality DNA templates where little or no heterogeneity is observed in the poly(A/T) sequence, the resulting mRNA exhibited additional heterogeneity in the poly(A) tail, primarily resulting from transcriptional slippage of T7 RNA polymerase, resulting in non-templated additional adenosine nucleotides.

Further characterisation of the mRNA poly(A) tail length and heterogeneity from a wide range of alternative linearised DNA templates demonstrated that the presence of additional nucleotides downstream of the poly(T) sequence in the linearised DNA template reduced transcriptional slippage of T7 polymerase and therefore reduced mRNA poly(A) tail heterogeneity. Moreover, the analysis of RNase T1-digested mRNA using LC-MS enabled the analysis of the 3’-end of the mRNA and any potential heterogeneity arising through 3’-end additions of the mRNA generated from the DNA templates that are linearised within the poly(T) sequence. The LC-MS analysis revealed that multiple populations were observed from mRNA generated from these DNA templates, consistent with the 3’-end addition of cytidine.

Furthermore, in this study, we demonstrate how manufacturing conditions such as NTP concentration impact the mRNA poly(A) tail length heterogeneity. Using lower ATP concentrations resulted in a decrease in mRNA poly(A) tail heterogeneity by reducing T7 RNA polymerase transcriptional slippage in the mRNA generated from DNA templates linearised within the poly(T) DNA template. In addition, 3’-end heterogeneity was also increased when CTP concentration was increased, further validating previous observations. No significant differences in mRNA poly(A) tail length heterogeneity were observed during the time course of an IVT reaction.

This work demonstrates the power of LC-MS analysis of DNA templates using restriction digestion and direct LC-MS analysis to rapidly assess DNA template quality. This is an additional tool for identity testing of the DNA template, providing nucleotide resolution of the poly(A/T) sequence of DNA plasmids used for manufacturing mRNA medicines. Moreover, the impact of DNA template quality and manufacturing conditions provides further insight into the parameters that impact mRNA poly(A) tail length and heterogeneity, an important critical quality attribute. The ability to rapidly assess DNA template quality, combined with monitoring and controlling mRNA poly(A) tail length and heterogeneity, will ensure the consistent quality of mRNA from manufacturing processes and provide important information as part of detailed characterisation studies of precision mRNA medicines. Such information will be important for establishing acceptance criteria and the transfer of methods to quality control and release assays.

## MATERIALS AND METHODS

### Restriction digestion and linearisation of plasmid DNA (preparation of plasmid DNA)

Linearisation of plasmid DNA in preparation for IVT was performed using restriction enzymes sourced from either New England Biolabs or Fisher Scientific. Approximately 30-50 µg of plasmid DNA was digested using 100 units of restriction enzyme following the manufacturer’s instructions. Digestions were incubated overnight at 37 °C prior to agarose gel electrophoresis. DNA fragment purification was performed using the Monarch PCR & DNA Cleanup Kit (New England Biolabs) or by phenol-chloroform extraction (UltraPure™ Phenol:Chloroform:Isoamyl Alcohol, 25:24:1, v/v, Thermo Fisher Scientific). Restriction digestion for the analysis of the poly(A/T) region was performed as above with approximately 5 μg of DNA. Digestions with SfiI were performed at 50°C according to the manufacturer’s instructions.

### PCR amplification of DNA templates

PCR amplification was performed using the KAPA2G ReadyMix PCR kit (Roche) using primers upstream of the T7 promoter region and downstream of the poly(A/T) tail. Reaction mixes were prepared according to the manufacturer’s instructions to a final volume of 25 μL in nuclease-free water with final concentrations of 0.5 μM of each primer, 1 ng of DNA template, and 1X final concentration of KAPA2G master mix. Cycling conditions were performed as follows: Initial denaturation for 3 minutes at 95°C, followed by 30 cycles of denaturation for 30 seconds at 95°C, annealing for 15 seconds at 60°C, and extension for 15 seconds at 75°C. Final extension was 1 minute at 75°C. The PCR product was purified using the Monarch PCR & DNA Cleanup Kit (New England Biolabs), and the purified DNA was used directly as the template for *in vitro* transcription. The PCR product was digested with BsiWI, which cuts upstream of the poly(A/T) region, before purification and LC-MS analysis of the DNA template.

### Molecular cloning of a plasmid DNA template

In-house generation of a plasmid DNA template (4) IVT was performed using a commercially available cloning kit (Cloning Kit for mRNA Template, Takara) according to the manufacturer’s instructions. Briefly, the eGFP CDS fragment was amplified by PCR using primers containing the 15-base sequences for In-Fusion cloning into a pre-linearised vector with a T7 promoter, transcription start sequence (AGG),5’-UTR 3’-UTR, and a 105-base poly(A) sequence. All other linearised plasmid DNA templates (1, 2, 3, 5 and 6) were obtained from commercial suppliers.

### Sanger sequencing of DNA templates

Sanger sequencing of the homopolymeric poly(A/T) region of the DNA templates was performed using primers upstream and downstream of the poly(A/T) region on the plasmid template. Sequencing data was provided from the commercial DNA plasmid manufacturer and additional Sanger Sequencing was generated from GENEWIZ (Azenta Life Sciences).

### *In vitro* transcription

mRNA transcripts were prepared using IVT. DNA templates used in this study are summarised in Table 1. The general IVT composition included T7 bacteriophage DNA-dependent RNA polymerase (1.00 µM; Roche), and equimolar concentrations (10 mM each) of ATP, CTP, GTP, and UTP (Roche). The reaction mixture also contained magnesium acetate (42 mM), HEPES buffer (40 mM, pH 7; Thermo Fisher Scientific, USA), dithiothreitol (DTT, 10 mM), sodium chloride (50 mM), and spermidine (2 mM), all obtained from Sigma-Aldrich. Inorganic pyrophosphatase was added (3.00 µM, Roche) to prevent magnesium pyrophosphate precipitation, and RNase inhibitor (0.19 µM, Roche) was added to maintain RNase-free conditions. The reaction also contained 0.01% (v/v) Triton X-100 (Merck). Template DNA was present at 0.05 µg/µL. Reactions were incubated at 37 °C for two hours (unless specified otherwise), then quenched by adding EDTA (Thermo Fisher Scientific) to a final concentration of 80 mM. Following IVT, RNA was purified by solid-phase extraction using silica columns as previously described (37). RNA concentrations were measured using a NanoDrop™ OneC spectrophotometer (Thermo Fisher Scientific) by absorbance at 260 nm, normalised to a 1.0 cm (10.0 mm) path length. In select experiments, ATP concentrations were varied from 0.5 to 15 mM to assess their impact on poly(A) tail heterogeneity, while all other components were kept constant. Similar tests were conducted by varying UTP and CTP, either independently or in combination with ATP, using the same IVT conditions. Samples collected for time-course studies were quenched with EDTA as previously described.

### Poly(A) tail sample preparation

RNase T1 digests used to generate poly(A) tail containing fragments were performed in solution using 25-50 µg of mRNA with the addition of 60 U RNase T1 (Thermo Fisher Scientific) at 37 °C for 4 hrs or overnight in 0.1 M triethylammonium acetate (Sigma-Aldrich). 10 mM EDTA-free acid (Sigma-Aldrich) (adjusted to pH 8 with ammonium hydroxide, Sigma-Aldrich) was added prior to LC-MS to minimise metal cation adduction.

### Liquid Chromatography-Mass Spectrometry Analysis

IP-RP HPLC-UV/MS analysis was performed on a Vanquish UHPLC system (Thermo Fisher Scientific) online to an Orbitrap Exploris 240 mass spectrometer (Thermo Fisher Scientific). Separations were performed using a 250 mm x 2.1 mm DNAPac RP column (Thermo Fisher Scientific) and a C18 ACQUITY Premier Oligonucleotide C18 Column, 130Å, 1.7 µm, 2.1 x 100 mm column (Waters). LC-MS mobile phases were prepared in UHPLC-MS Grade water (Thermo Scientific). Mobile phase A was comprised of 10 mM dibutylamine (DBA, >99.0% (GC), Sigma-Aldrich) and 50 mM 1,1,1,3,3,3-hexafluoro-2-propanol (HFIP, 99.5+%, Thermo Scientific ). Mobile phase B was comprised of 10 mM DBA and 50 mM HFIP with 50% acetonitrile (UHPLC-MS grade, Thermo Scientific) at 0.25 mL min^-1,^ column temperature 50 °C with UV detection at 260 nm.

LC separation of plasmid restriction digests was performed starting at 30% B with a linear extension over 1 minute to 35% B and increased to 52% over 19 mins (curve 3). For analysis of mRNA poly(A) tails, the gradient started at 35% B with a linear extension over 1 minute to 40% B to 50% over 30 minutes. Mass spectrometry data acquisition was performed using an Orbitrap Exploris 240 instrument with a heated-electrospray ionisation source with spray voltage of 2500V in negative mode. Source settings were sheath gas 35 au, aux gas 10 au, ion transfer tube temperature 320°C, and vaporiser temperature 350°C. Data was acquired in ‘Profile’ mode using the ‘Intact Protein’ application mode and the ‘Low Pressure’ mode. For DNA template analysis, MS1 spectra were captured at Orbitrap resolutions of 15,000 and 120,000 with a scan range of 450-2500 m/z, RF lens 75%, a normalised AGC target of 100%, and a 100 ms maximum injection time. 3 microscans were performed in negative mode. For the analysis of poly(A) tails, MS1 spectra were captured at an Orbitrap resolution of 240,000 with a scan range of 750-5500 m/z, RF lens 75%, a normalised AGC target of 100%, and a 300 ms maximum injection time. 5 microscans were performed in negative mode.

### LC-MS data analysis

LC-MS data was analysed using BioPharma Finder 5.2 (Thermo Fisher Scientific). using the ‘Intact Mass Analysis’ experiment type. Isotopically unresolved LC-MS data acquired at 15,000 resolution was processed using the ReSpect™ deconvolution algorithm whereas the isotopically resolved 240,000 resolution data was analysed using the Xtract™ deconvolution algorithm. Mass deconvolution employed the ‘Sliding Windows’ ‘Source Spectra’ method. Sliding window parameters were adjusted to accommodate the chromatographic peaks. The ‘Peak Model’ was set to ‘Nucleotide’ with a ‘Negative Charge’ specified for deconvolution.

Deconvolution and annotation of mRNA poly(A) tails was performed using the ‘Intact Protein Deconvolution’ function in Chromeleon 7.3.1 using the Xtract™ deconvolution algorithm using the ‘Sliding Windows’ ‘Source Spectra’ method as previously described. Custom columns, Target Mass (Da) and Target Tolerance (Da), were added to the sequence manager window. Target Mass was set to the mass of the most abundant tail length, and Target Tolerance was set to 1. A custom impurities table featuring the ions specific to this analysis was used with the reporting feature to quantify the target mass, and the mass of n-x and n+x poly(A) tails relating to the mRNA sequence. For LC-MS analysis replicate data shown in Supplementary Figures S7-25.

## DATA AVAILABILITY

The mass spectrometry data have been deposited to the ProteomeXchange Consortium via the PRIDE partner repository with the dataset identifier PXD071020.

Project accession: PXD071020 Token: dfmk3xUZbGxs

Alternatively, reviewer can access the dataset by logging in to the PRIDE website using the following account details:

Username

Password: BPpObaBFJwb9

## ACKNOWLEDGEMENTS

The authors acknowledge Rebecca Ryan for procuring the NLuc plasmid DNA.

Graphical abstract created in BioRender. Dickman, M. (2025) https://BioRender.com/6nysa7h

## AUTHOR CONTRIBUTIONS

Gareth Owen: Conceptualisation, Formal analysis, Investigation, Methodology, Visualisation, Writing – original draft, Writing – review and editing. Caroline Evans: Conceptualisation, Formal analysis, Investigation, Methodology. Adithya Nair: Conceptualisation, Investigation, Methodology. Sebastian Ross: Conceptualisation, Investigation, Methodology. Mollie Glenister: Formal analysis, Visualisation. Zoltán Kis: Funding acquisition, Supervision, Writing – Review & Editing. Mark Dickman: Conceptualisation, Funding acquisition, Project administration, Supervision, Writing – original draft, Writing - review & editing.

## SUPPLEMENTARY DATA

Supplementary Data are available at NAR online.

## DATA AVAILABILITY

Project accession: PXD071020 Token: dfmk3xUZbGxs

Username

Password: BPpObaBFJwb9

