## Supplementary Figures for "DNA template heterogeneity and *in vitro* transcription reaction conditions impact the poly(A) tail length and heterogeneity of mRNA"

A

B

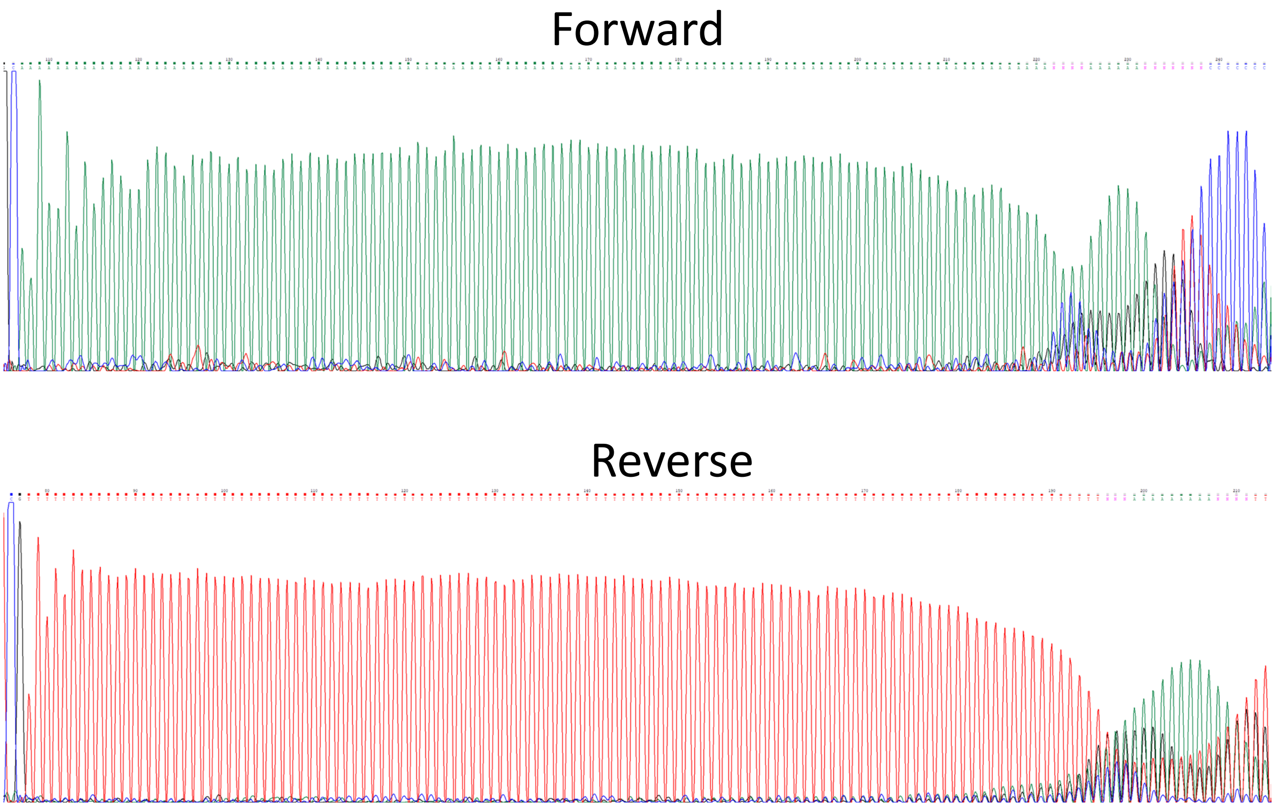


**Supplementary Figure S1. Sanger sequencing chromatograms covering the homopolymeric region of DNA template 1 encoding an NLuc mRNA.** (A) A representative forward read from the plasmid DNA template – 'A'-base reads shown in green (B). A representative reverse read from the plasmid DNA template – 'T'-base reads shown in red.

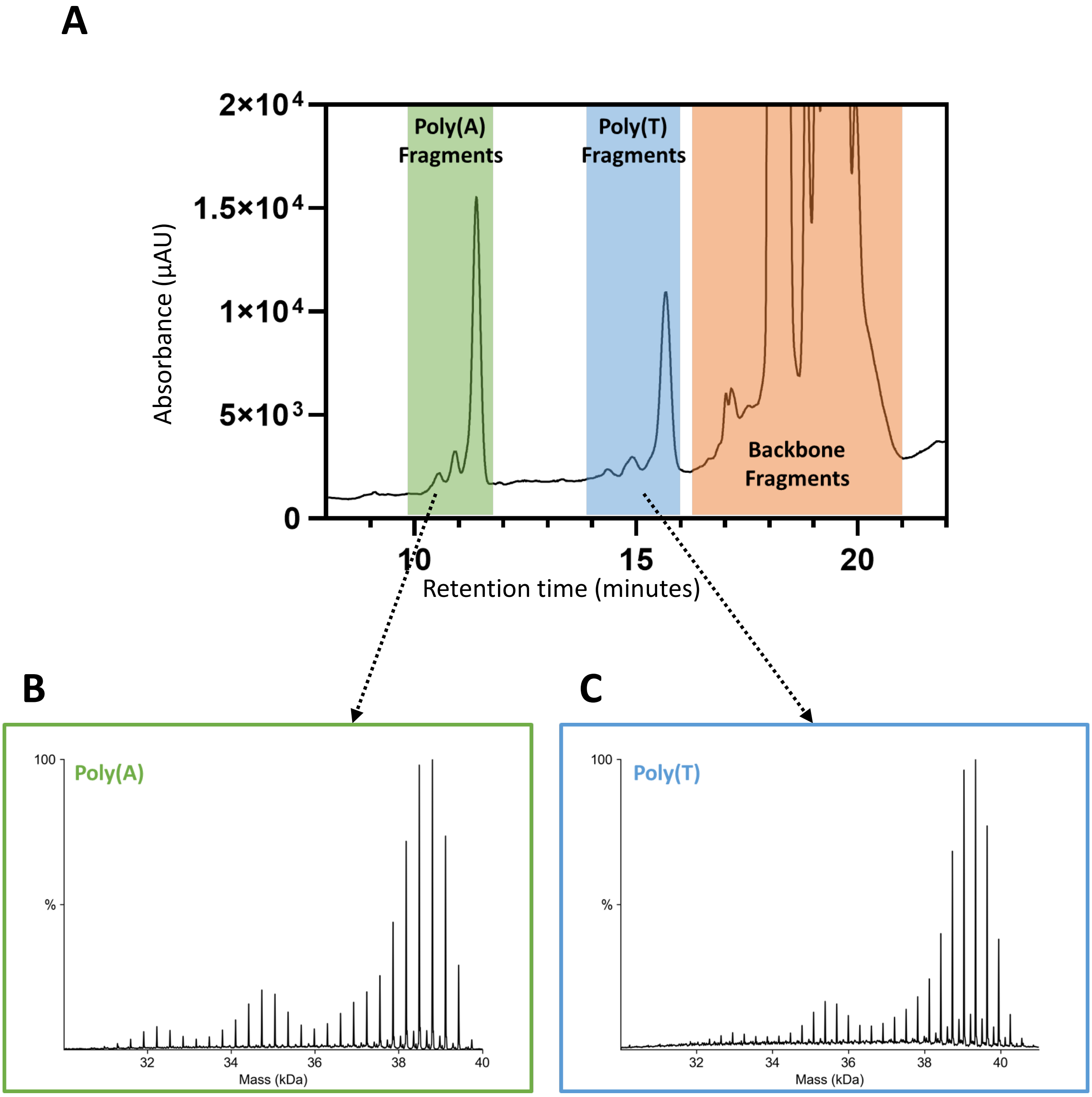


**Supplementary Figure S2. LC-UV and LC-MS analysis of a plasmid DNA template. (A) LC-UV chromatogram from a representative direct restriction digestion/LC-MS analysis of a DNA template encoding NLuc mRNA.** Chromatographic peaks corresponding to the poly(A)-containing sense strand species are highlighted in green and the peaks corresponding to the poly(T)-containing antisense strand species are highlighted in blue. Additional backbone restriction fragments are highlighted in orange. (B) Deconvoluted spectrum across the chromatographic peaks corresponding to the poly(A) containing species. (C) Deconvoluted spectrum across the chromatographic peaks corresponding to the poly(T) containing species.

### DNA Template 7 (eGFP)

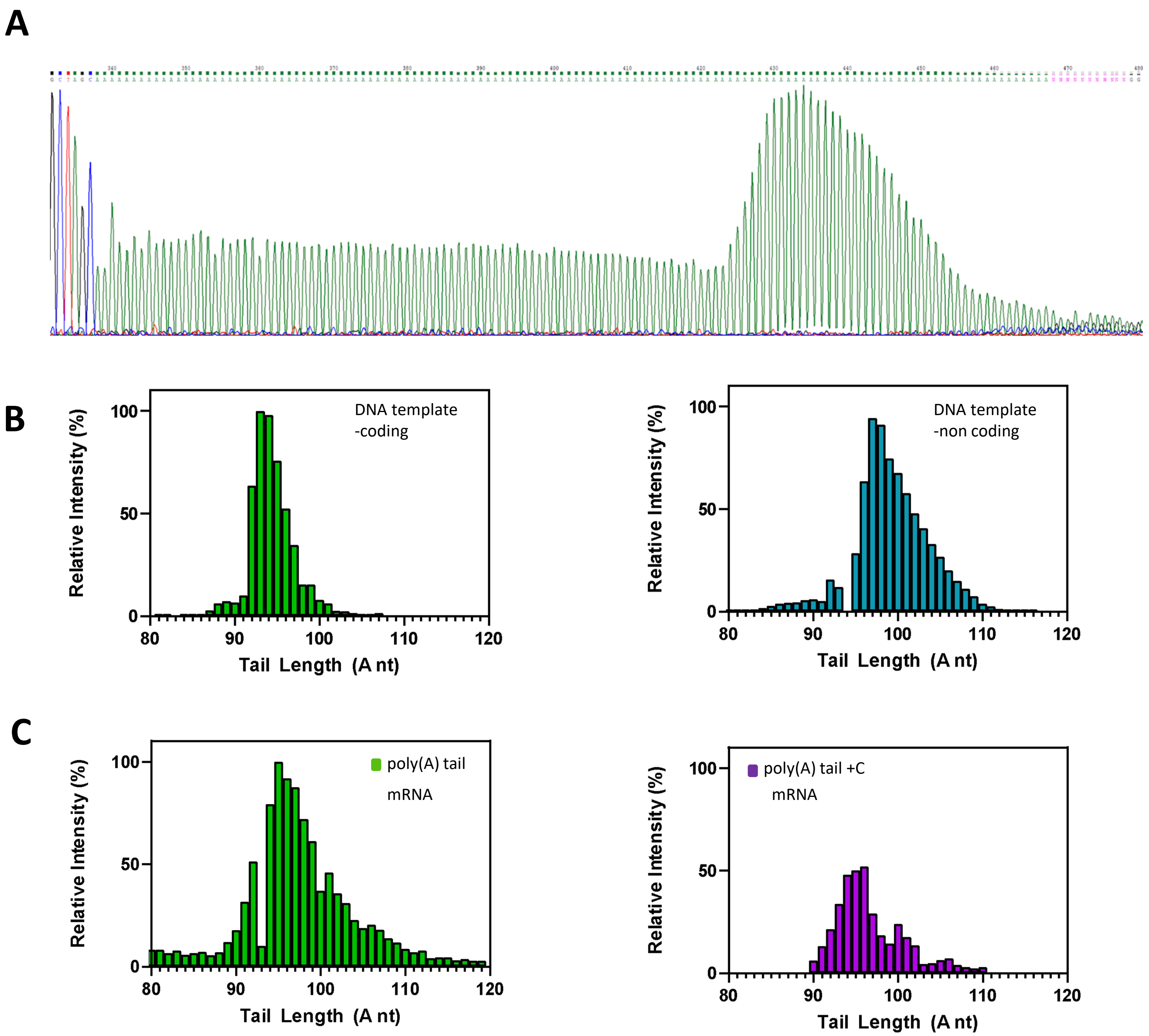


**Supplementary Figure S3. DNA template and mRNA poly(A) tail analysis of DNA template 7 encoding eGFP mRNA.** Analysis of an enzymatically produced DNA template and the resultant mRNA. (A) A representative Sanger sequencing chromatogram of the poly(A) encoding region of the DNA template used for *in vitro* transcription. (B) LC-MS identifications of tail species from the sense/poly(A) and antisense/poly(T) strands from restriction digestion of the synthetic DNA template. (C) mRNA poly(A) tail species identified by LC-MS from RNase T1 digestion of *in vitro* transcribed material. The poly(A) tail population is shown in green, and the poly(A)+C population is shown in purple.

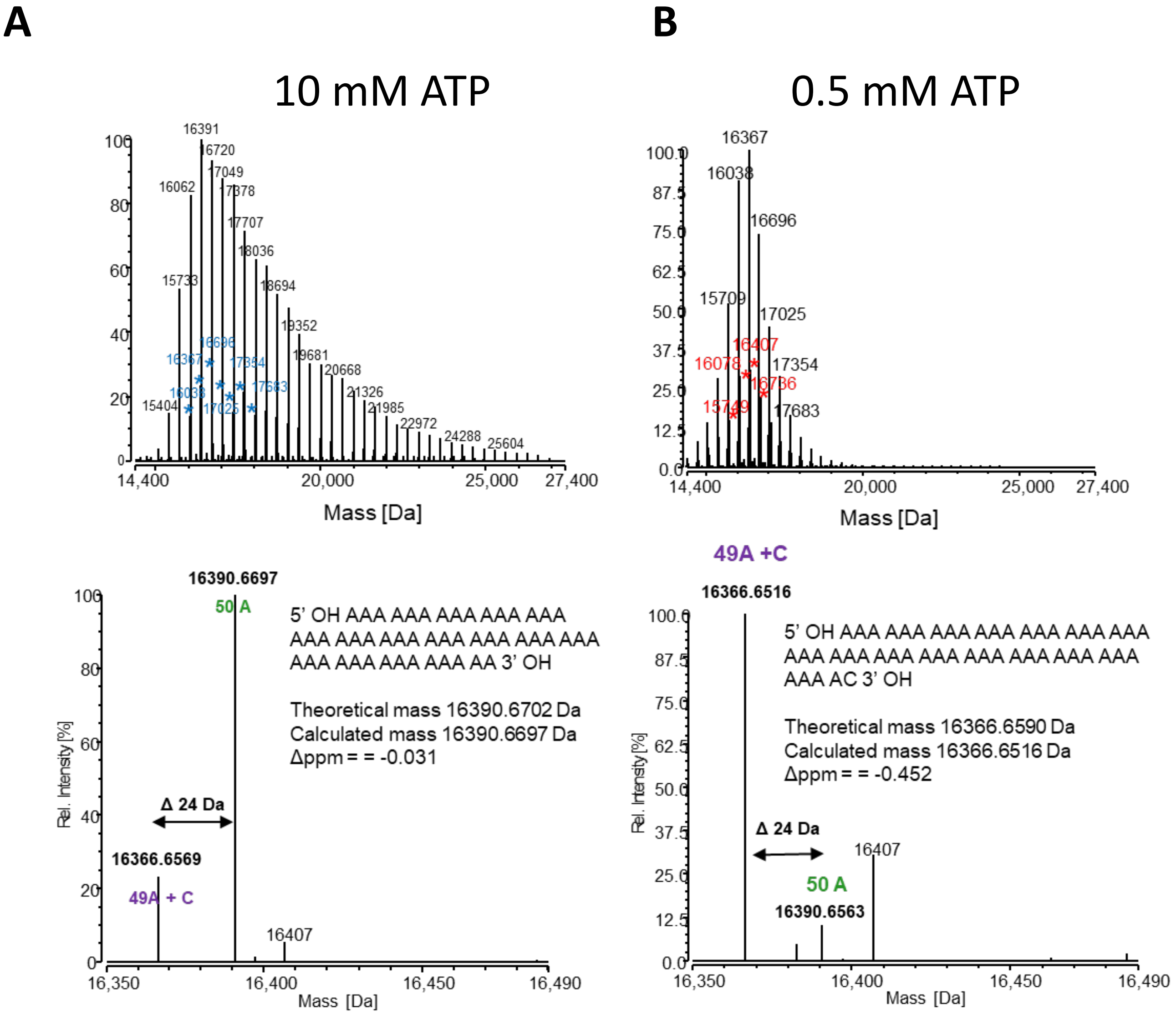


**Supplementary Figure S4. High-resolution accurate mass analysis of the two poly(A) tail distributions transcribed from DNA template 2**. Raw deconvoluted spectra from the LC-MS analysis of RNase T1 poly(A) tail fragments from mRNA transcribed from DNA template 2 using either 10 mM ATP (A) or 0.5 mM ATP (B). The theoretical and calculated masses are shown alongside the matched mass error. The expected 24 Da mass difference between 49A+C and 50A is shown.

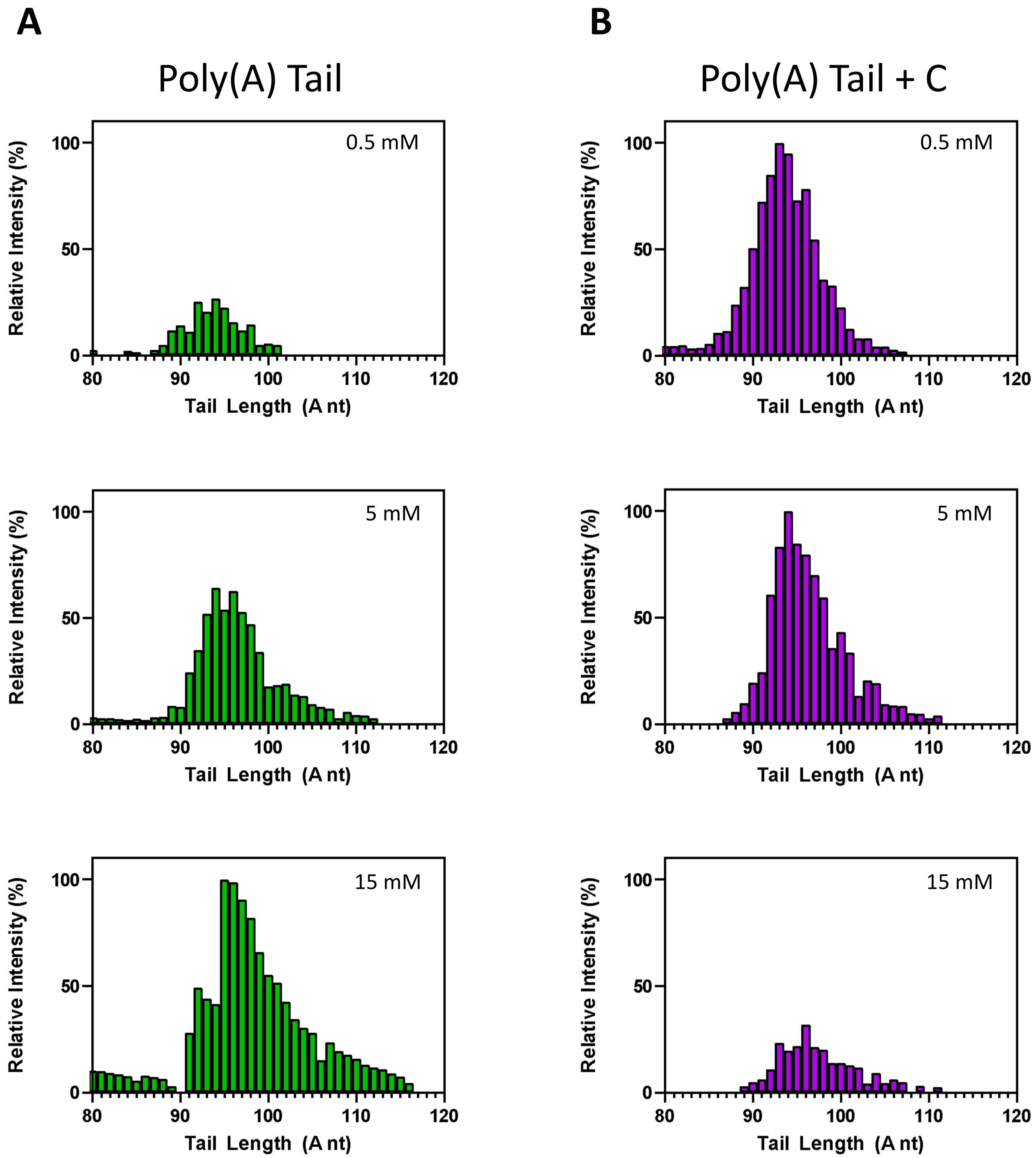


**Supplementary Figure S5. ATP Concentration Effects on the Poly(A) Tail Distribution of Template 7 (eGFP)** (**A**) mRNA poly(A) tail species identified by LC-MS from RNase T1 digestion of *in vitro* transcribed material using 0.5 mM, 5 mM, or 15 mM ATP. (**A**) The poly(A) tail population is shown in green. (**B**) The poly(A)+C tail population is shown in purple. The enzymatically produced DNA template does not contain additional bases after the poly(A) tail and is used directly in the IVT.

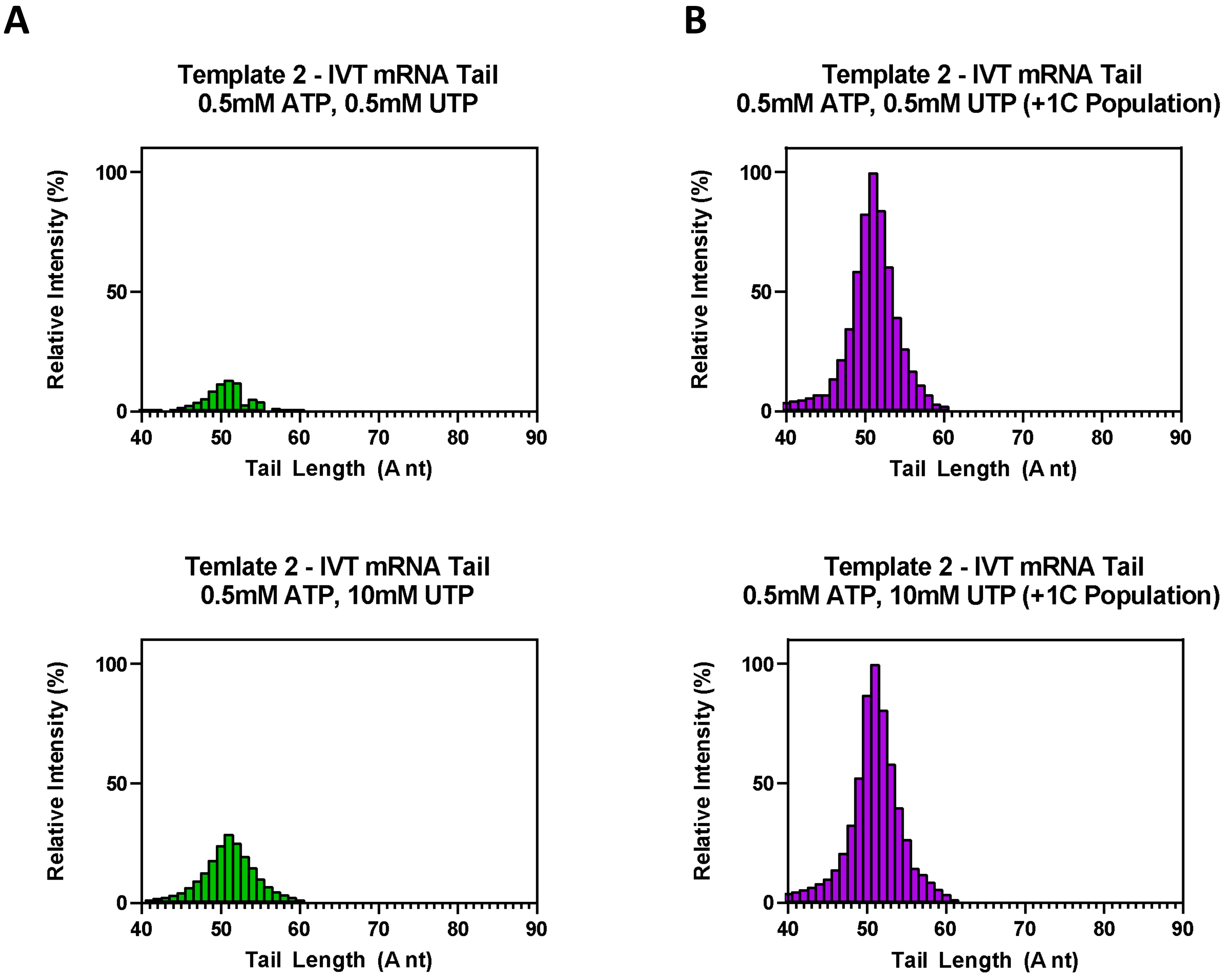


**Supplementary Figure S6. Effect of UTP concentration on the mRNA poly(A) tail length on heterogeneity**. mRNA was IVT from DNA template 2 (eGFP) and mRNA poly(A) tail species identified by LC-MS from RNase T1 digestion of *IVT* material produced using 0.5 mM ATP and either 0.5 mM or 10 mM UTP. (**A**) poly(A) tail population shown in green. (**B**) poly(A)+C tail population shown in purple.

### Replicate LC-MS Data
### Supplementary Figures S7-S25

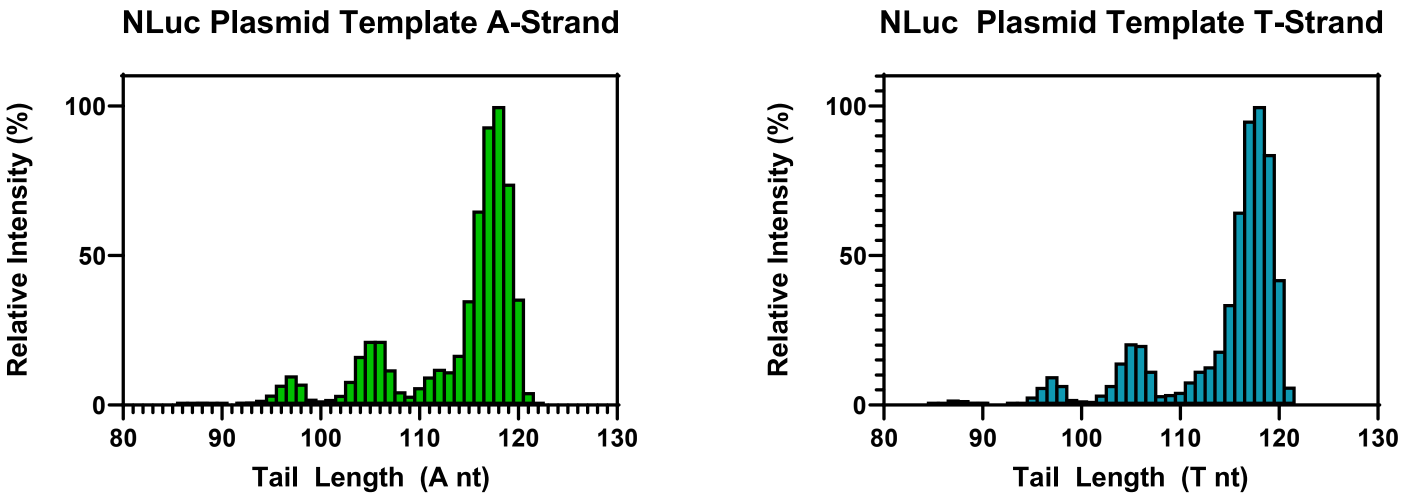


Template 1 – Rep. 1 – 120K resolution – Extended RT range

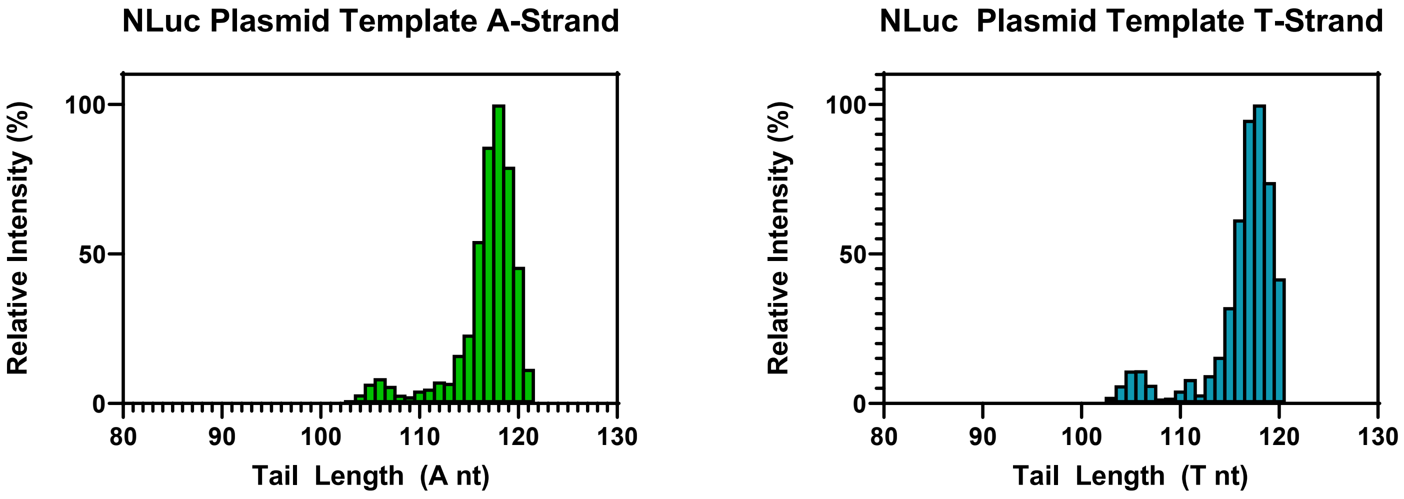


Template 1 – Rep. 2 – 15K resolution

**Supplementary Figure S7. Template 1 DNA LC-MS Analysis.** Tail identifications from LC-MS analysis of DNA template restriction fragments.

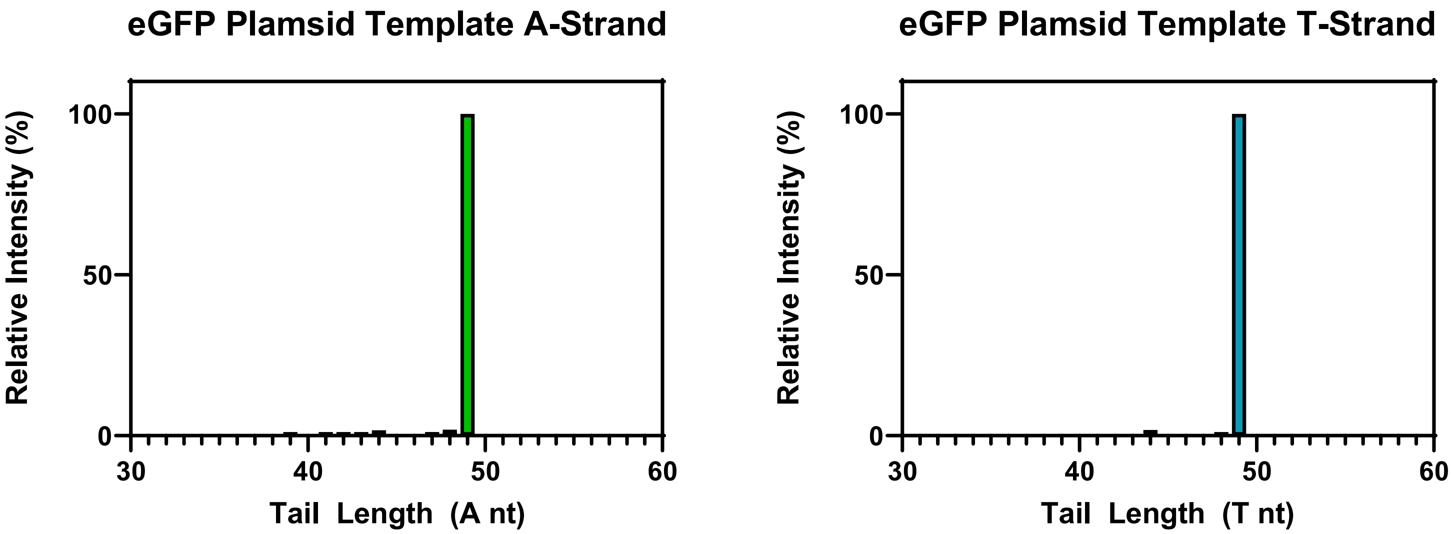


Template 2 – Rep. 1 – 120K resolution

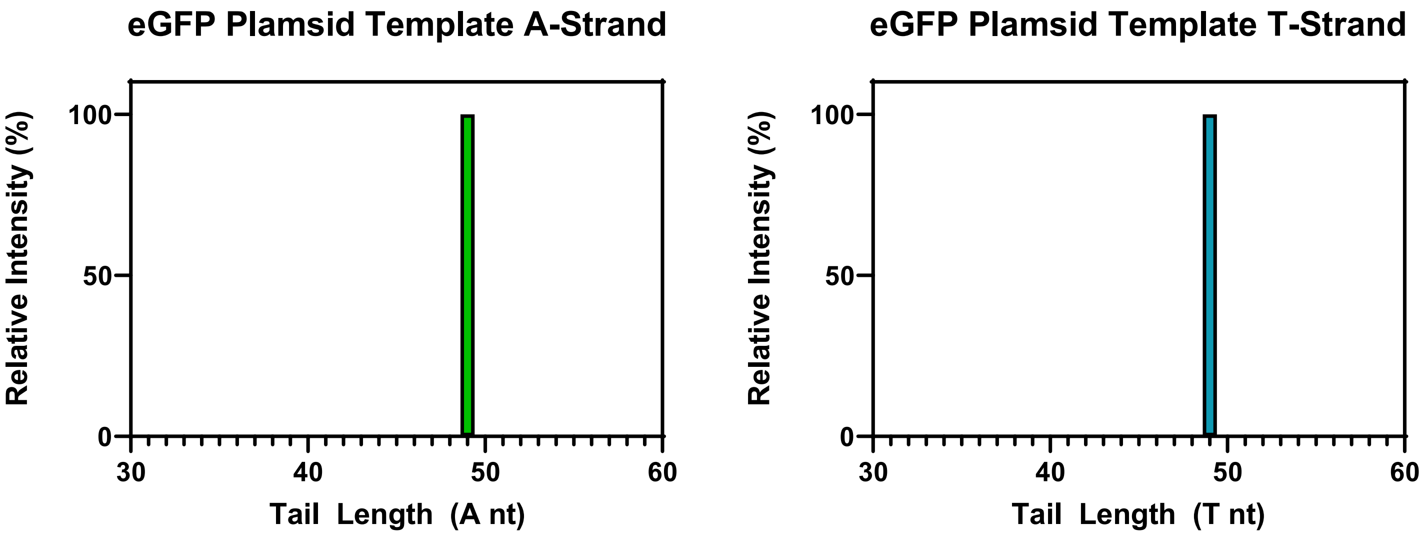


Template 2 – Rep. 2 – 15K resolution

**Supplementary Figure S8. Template 2 DNA LC-MS Analysis.** Tail identifications from LC-MS analysis of DNA template restriction fragments**.**

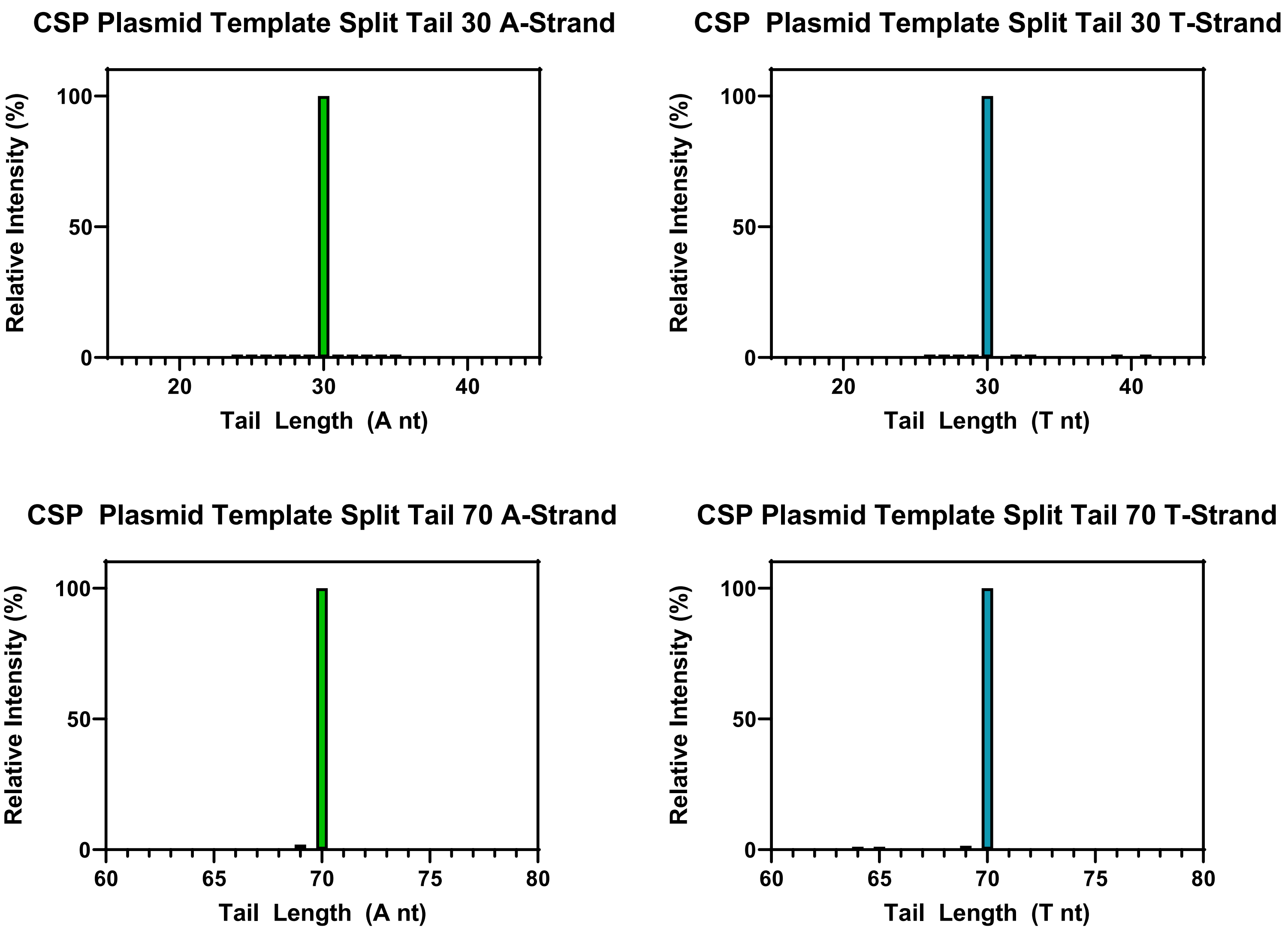


Template 3 – Rep. 1 – 120K resolution – Cut inside linker

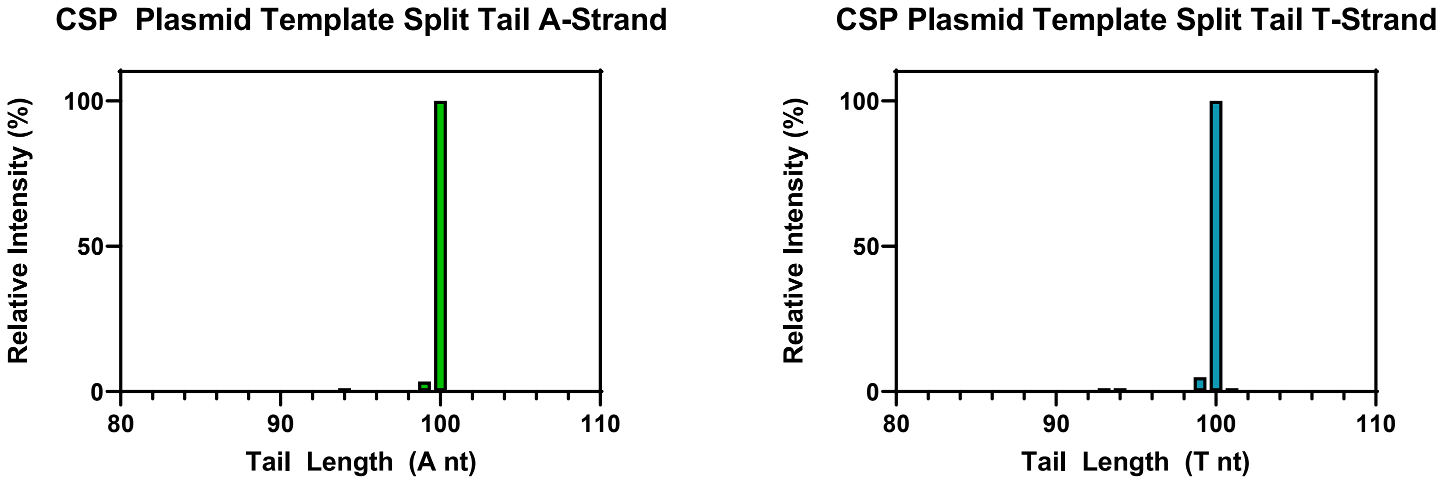


Template 3 – Rep. 2 – 15K resolution – Cut outside linker

**Supplementary Figure S9. Template 3 DNA LC-MS Analysis.** Tail identifications from LC-MS analysis of DNA template restriction fragments.

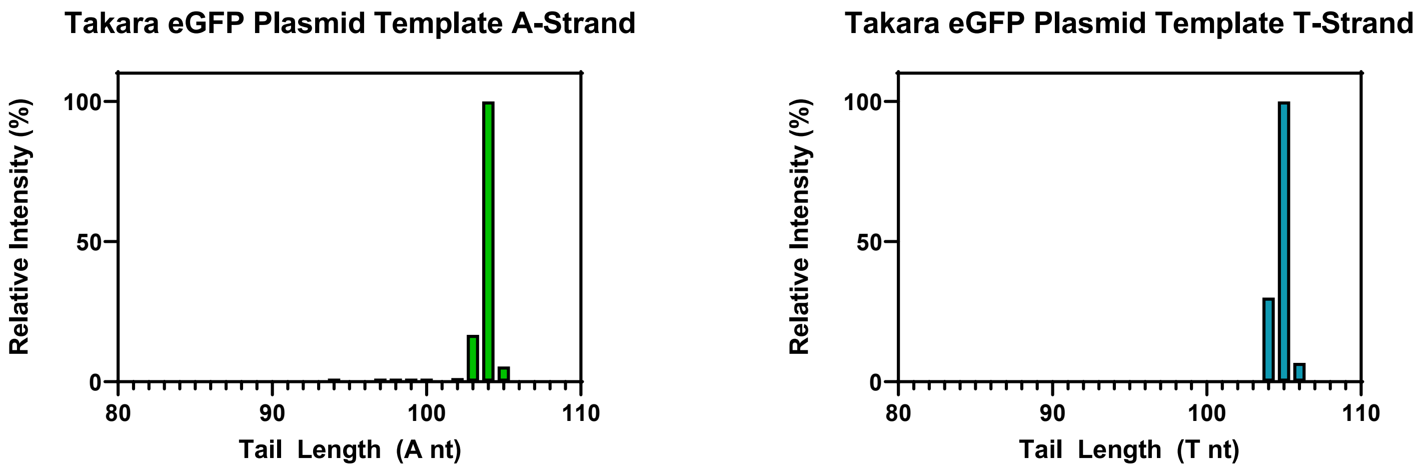


Template 4 – Rep. 1 – 120K resolution

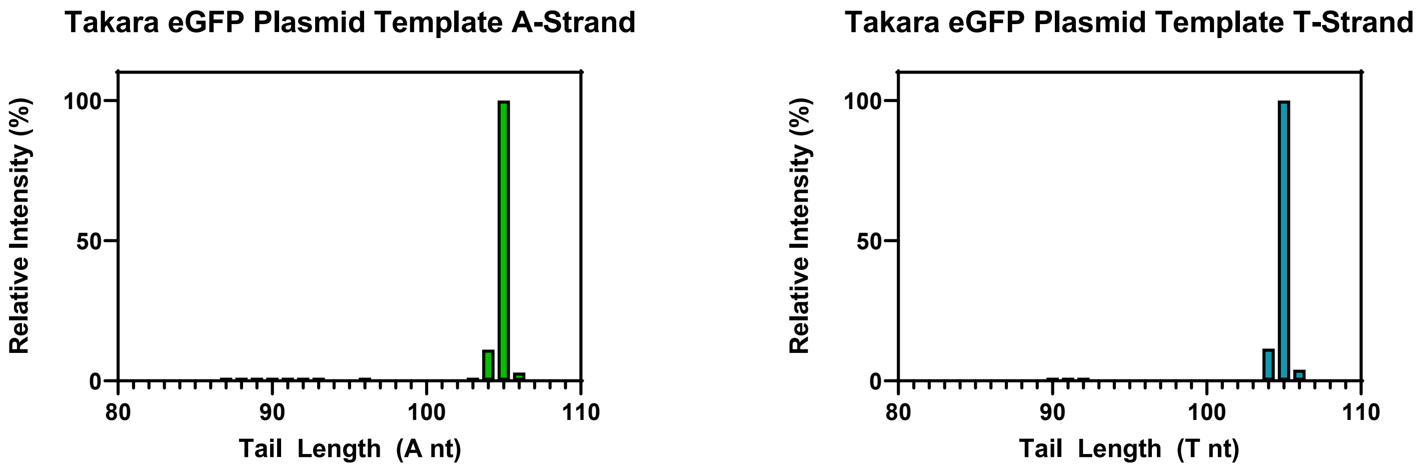


Template 4 – Rep. 2 – 15K resolution

**Supplementary Figure S10. Template 1 DNA LC-MS Analysis.** Tail identifications from LC-MS analysis of DNA template restriction fragments**.**

Rep. 1

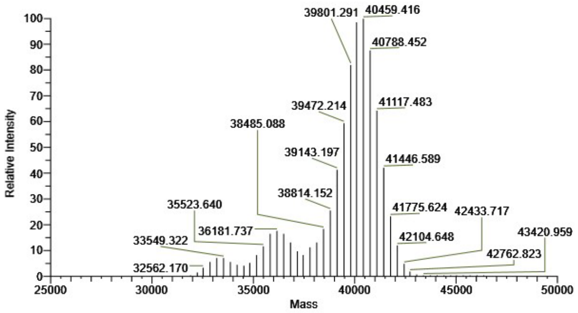


Rep. 2

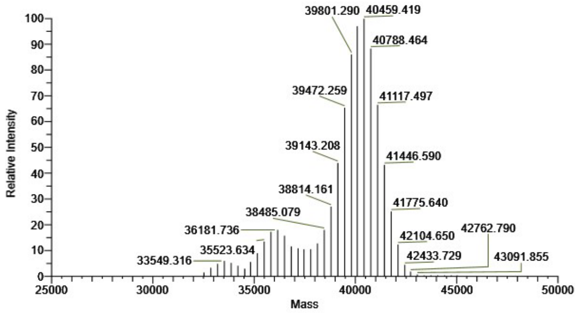


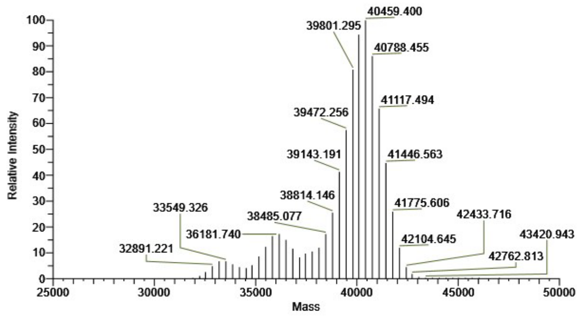


Rep. 3

**Supplementary Figure S11. Template 1 mRNA LC-MS Analysis.** Deconvoluted spectra of mRNA poly(A) tail species identified by LC-MS from RNase T1 digestion of *in vitro* transcribed material.

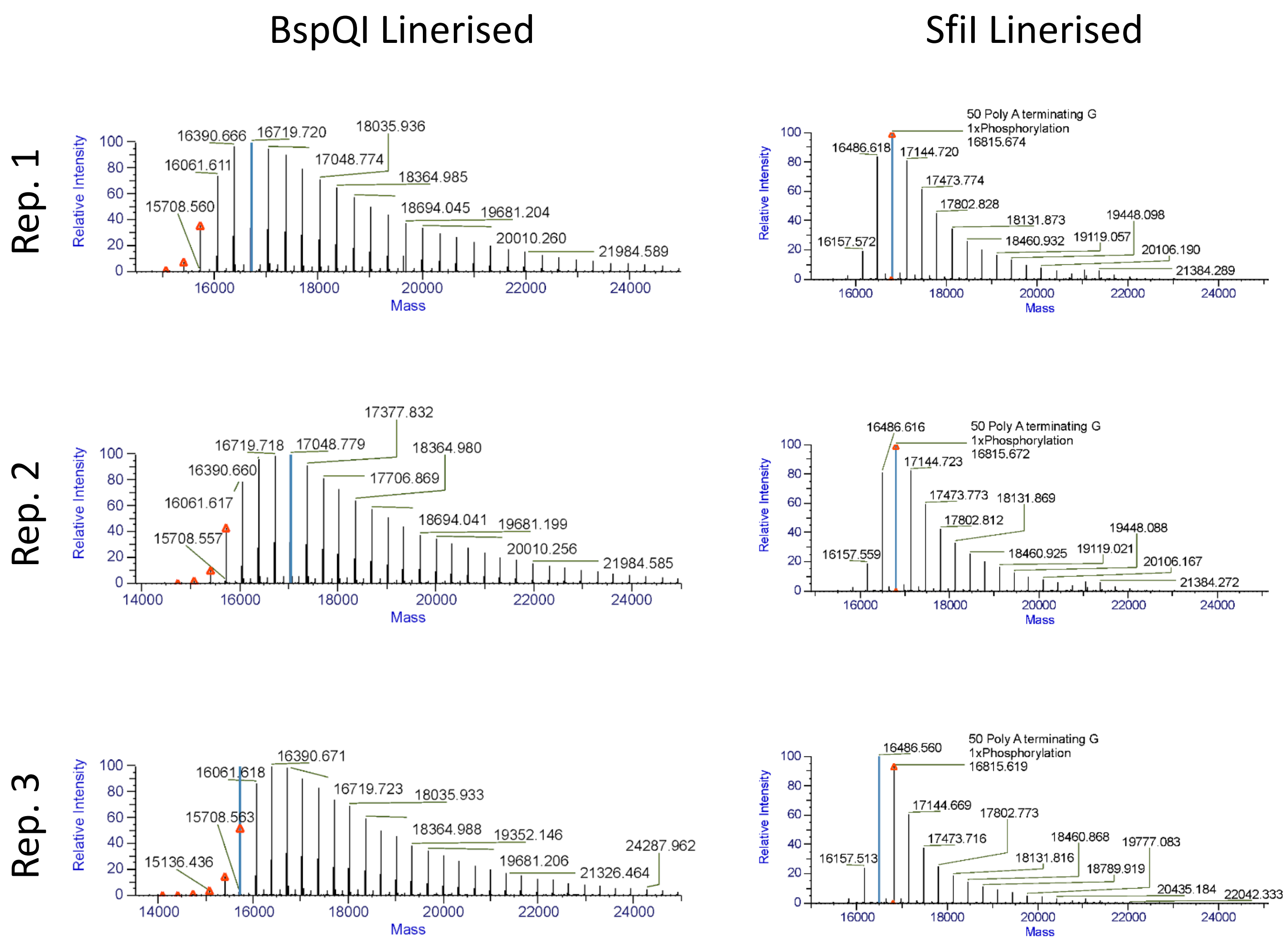


**Supplementary Figure S12. Template 2 mRNA LC-MS Analysis.** Deconvoluted spectra of mRNA poly(A) tail species identified by LC-MS from RNase T1 digestion of *in vitro* transcribed material from template linearised with either BspQI or SfiI.

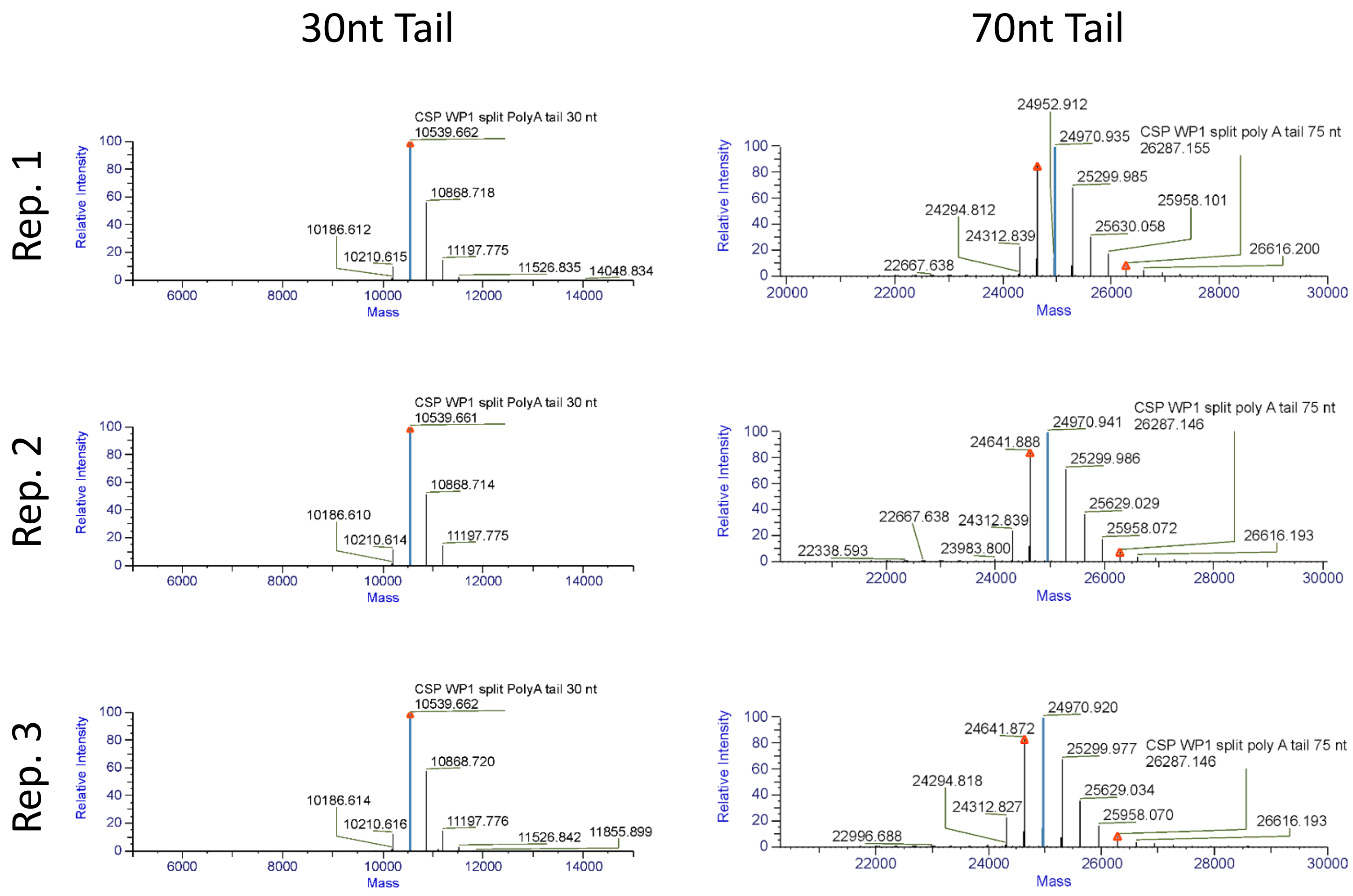


**Supplementary Figure S13. Template 3 mRNA LC-MS Analysis.** Deconvoluted spectra of mRNA poly(A) tail species identified by LC-MS from RNase T1 digestion of *in vitro* transcribed material. RNase T1 cleaves within the linker region of the split tail, generating two fragments with ~30As or ~70As.

Rep. 1

Rep. 2

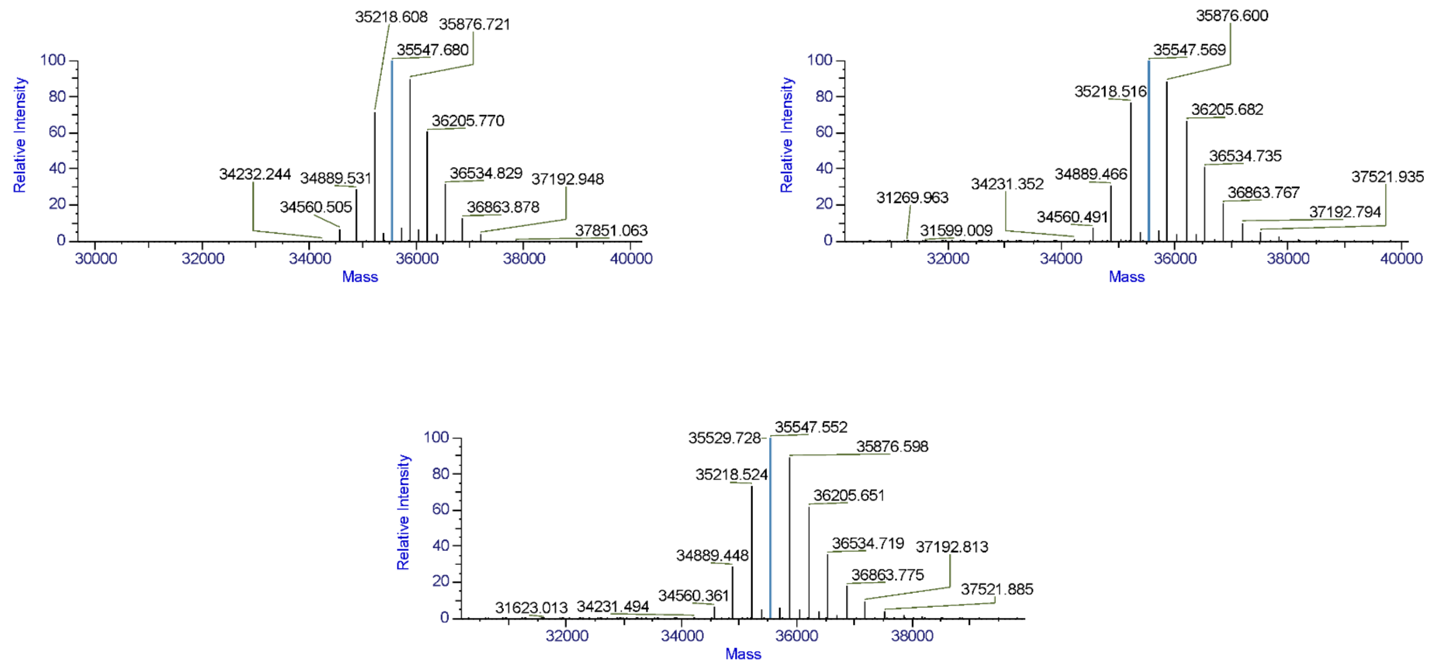


Rep. 3

**Supplementary Figure S14. Template 4 mRNA LC-MS Analysis.** Deconvoluted spectra of mRNA poly(A) tail species identified by LC-MS from RNase T1 digestion of *in vitro* transcribed material.

Rep. 1

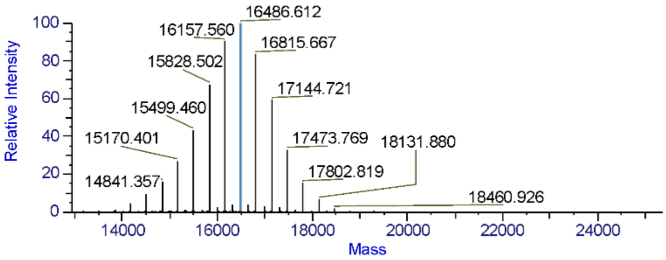


Rep. 2

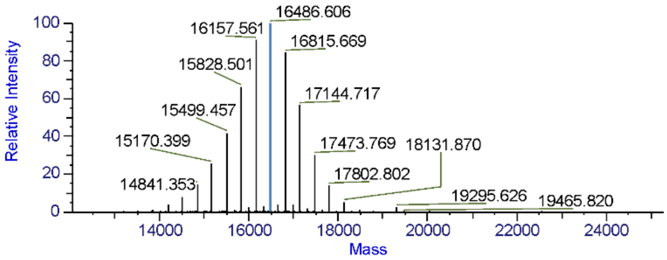


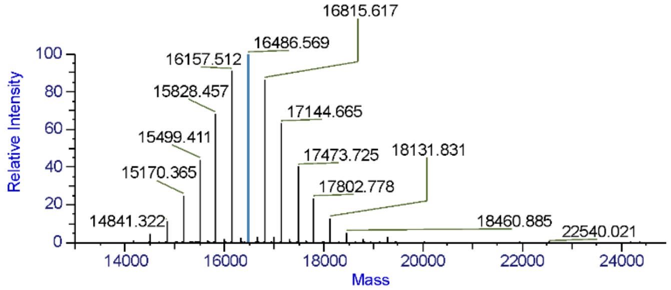


Rep. 3

**Supplementary Figure S15. Template 5 mRNA LC-MS Analysis.** Deconvoluted spectra of mRNA poly(A) tail species identified by LC-MS from RNase T1 digestion of *in vitro* transcribed material.

Rep. 1

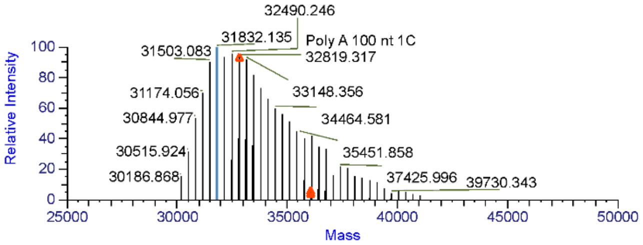


Rep. 2

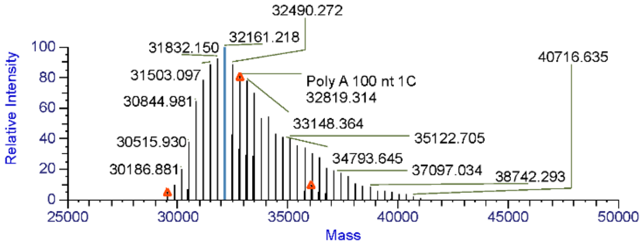


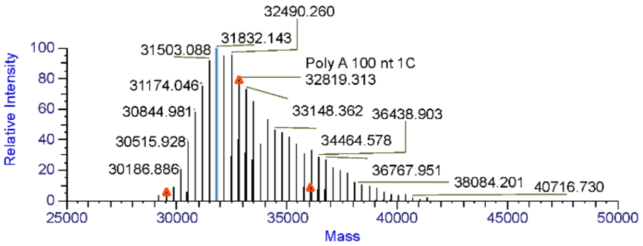


Rep. 3

**Supplementary Figure S16 – Template 6 mRNA LC-MS Analysis –** Deconvoluted spectra of mRNA poly(A) tail species identified by LC-MS from RNase T1 digestion of *in vitro* transcribed material.

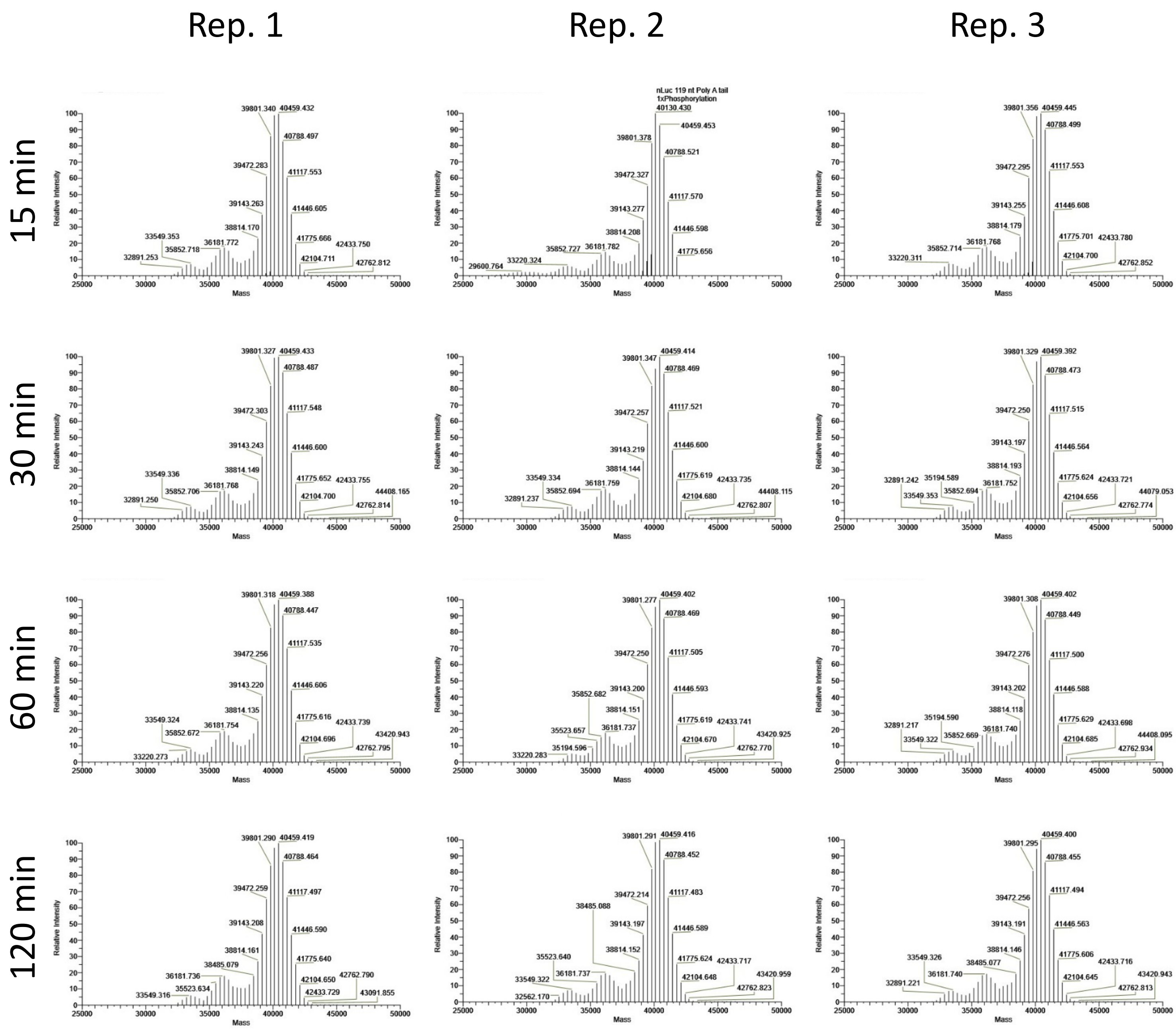


**Supplementary Figure S17. Template 1 mRNA IVT Time Course LC-MS Analysis.** Deconvoluted spectra of mRNA poly(A) tail species identified by LC-MS from RNase T1 digestion of *in vitro* transcribed material produced in IVT reaction of 15, 30, 60, or 120 minutes.

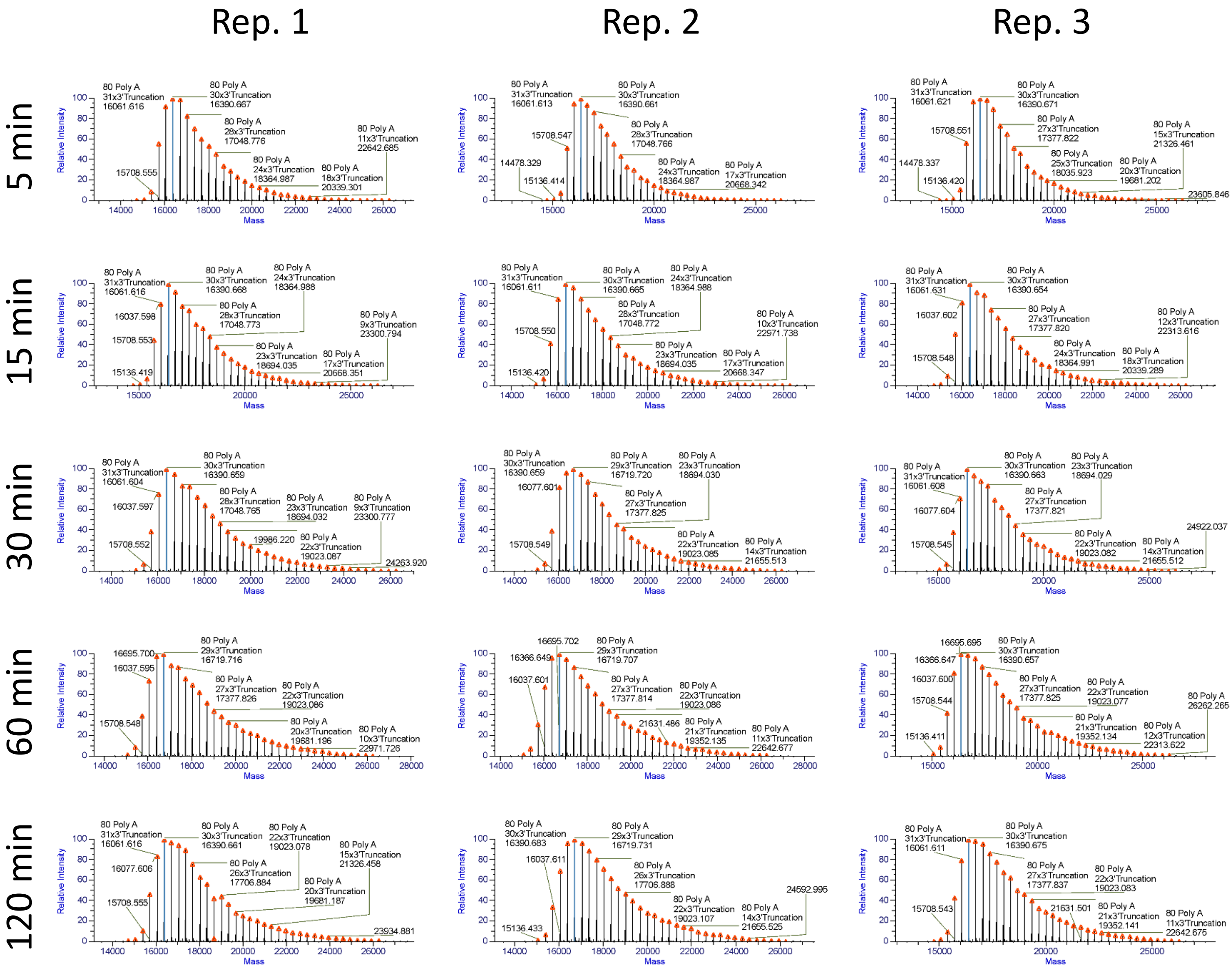


**Supplementary Figure S18. Template 2 mRNA IVT Time Course LC-MS Analysis.** Deconvoluted spectra of mRNA poly(A) tail species identified by LC-MS from RNase T1 digestion of *in vitro* transcribed material produced in IVT reaction of 15, 30, 60, or 120 minutes.

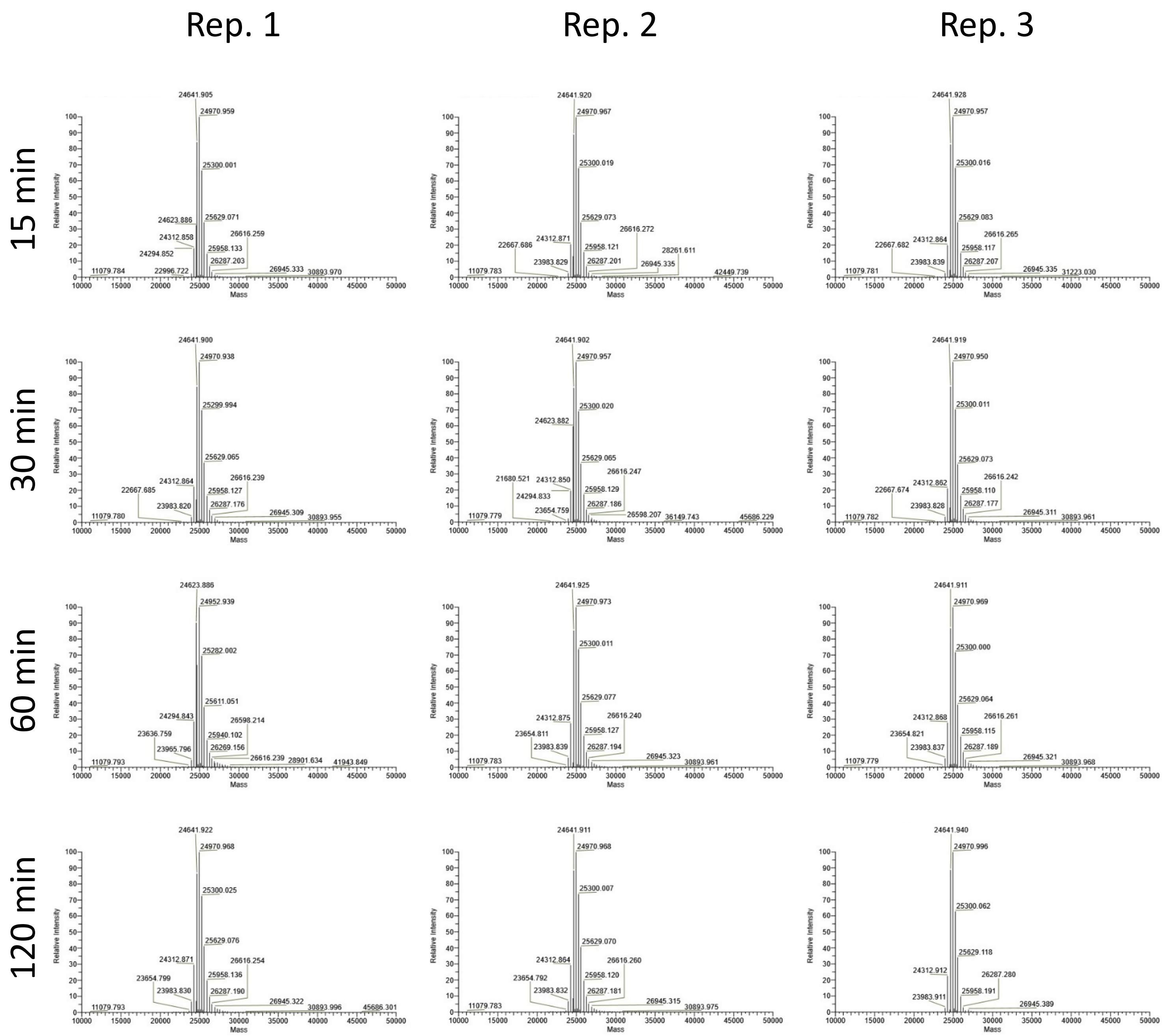


**Supplementary Figure S19. Template 3 mRNA IVT Time Course LC-MS Analysis.** Deconvoluted spectra of mRNA poly(A) tail species identified by LC-MS from RNase T1 digestion of *in vitro* transcribed material produced in IVT reaction of 15, 30, 60, or 120 minutes.

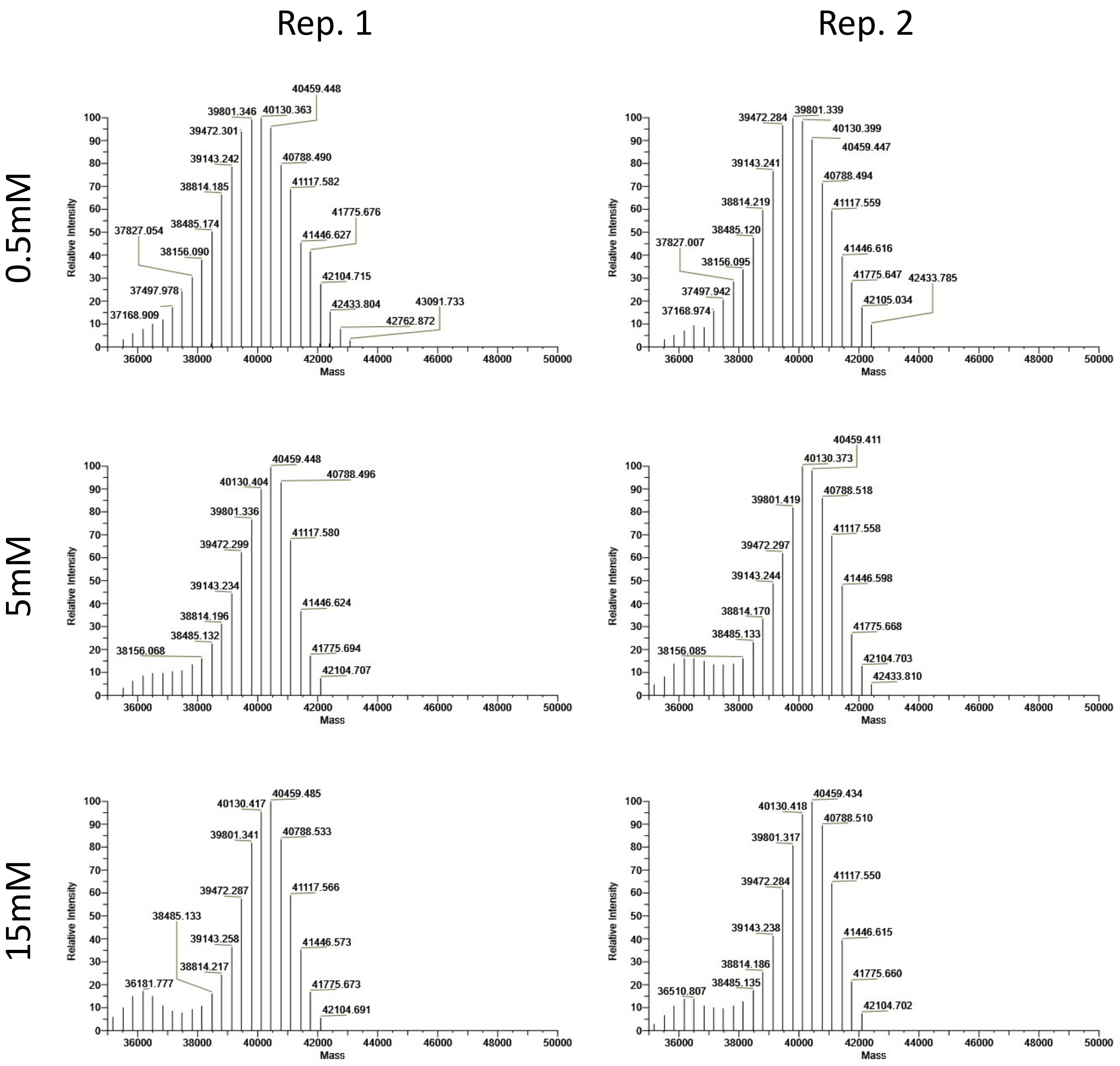


**Supplementary Figure S20. Template 1 mRNA ATP Concentration LC-MS Analysis.** Deconvoluted spectra of mRNA poly(A) tail species identified by LC-MS from RNase T1 digestion of *in vitro* transcribed material produced in IVT reaction with 0.5 mM, 5 mM, or 15 mM ATP.

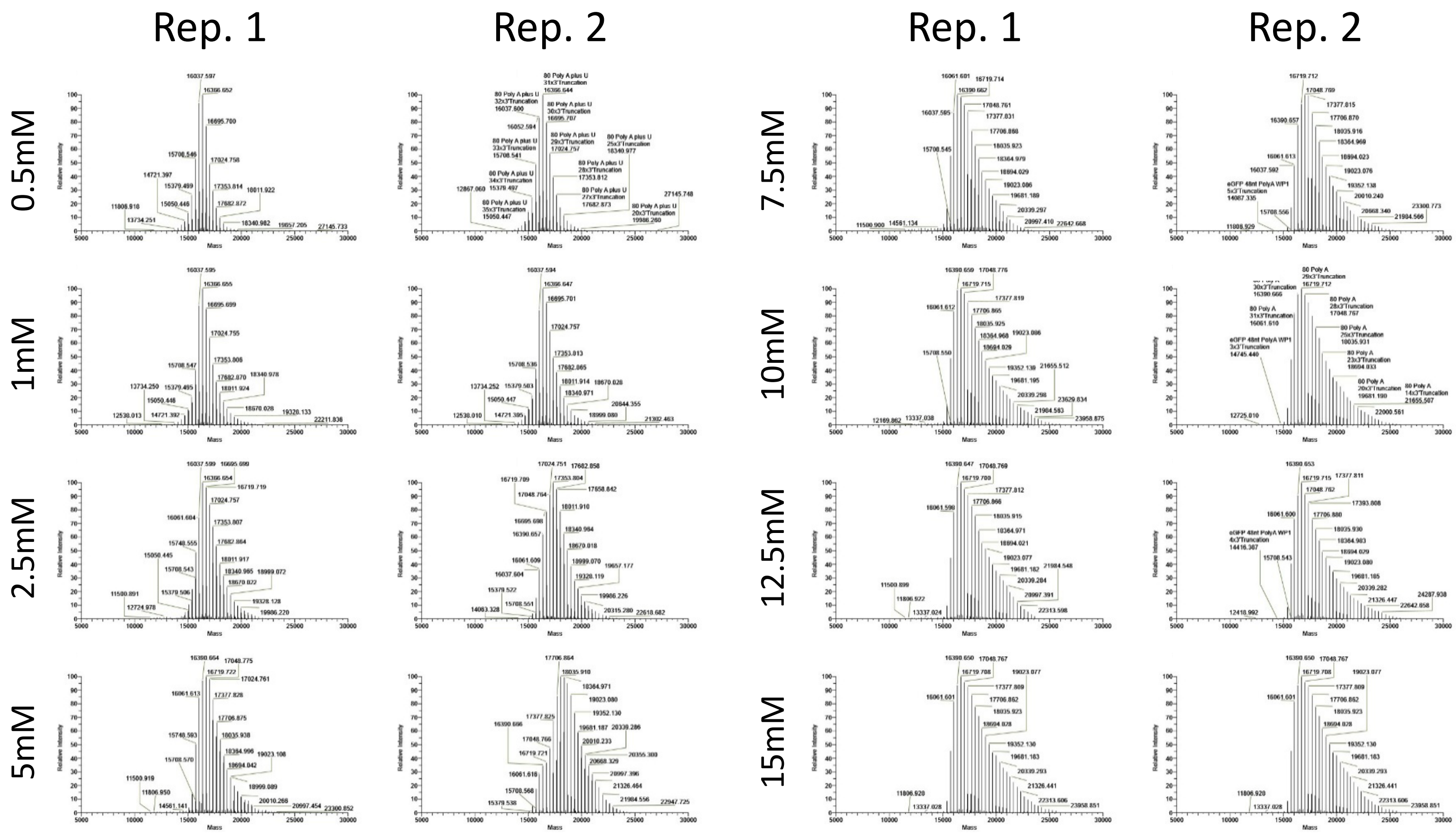


**Supplementary Figure S21. Template 2 mRNA ATP Concentration LC-MS Analysis.** Deconvoluted spectra of mRNA poly(A) tail species identified by LC-MS from RNase T1 digestion of *in vitro* transcribed material produced in IVT reaction with 0.5 mM, 1 mM, 2.5 mM, 5 mM, 7.5 mM, 10 mM, 12.5 mM or 15 mM ATP.

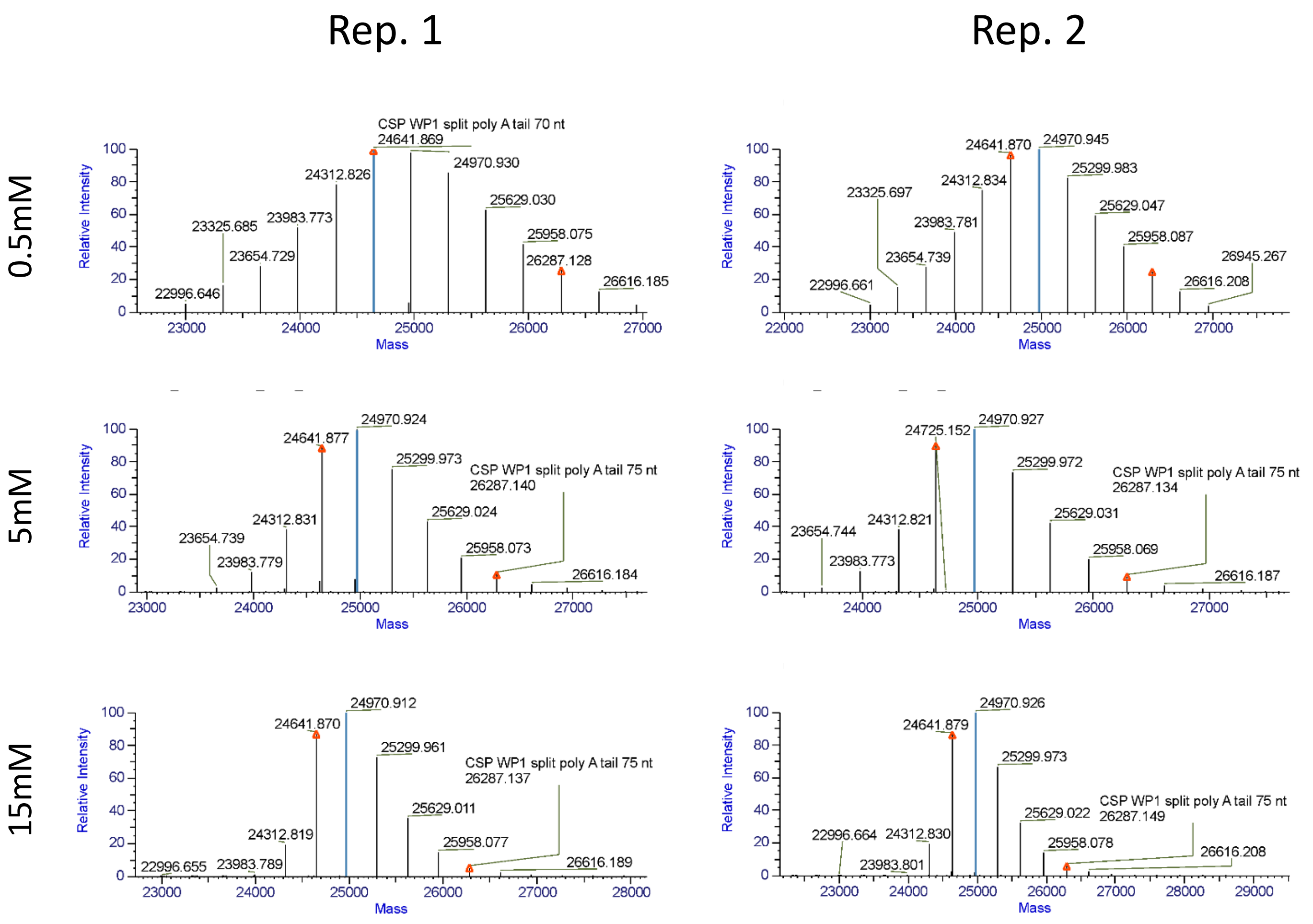


**Supplementary Figure S22. Template 3 mRNA ATP Concentration LC-MS Analysis.** Deconvoluted spectra of mRNA poly(A) tail species identified by LC-MS from RNase T1 digestion of *in vitro* transcribed material produced in IVT reaction with 0.5 mM, 5 mM, or 15 mM ATP.

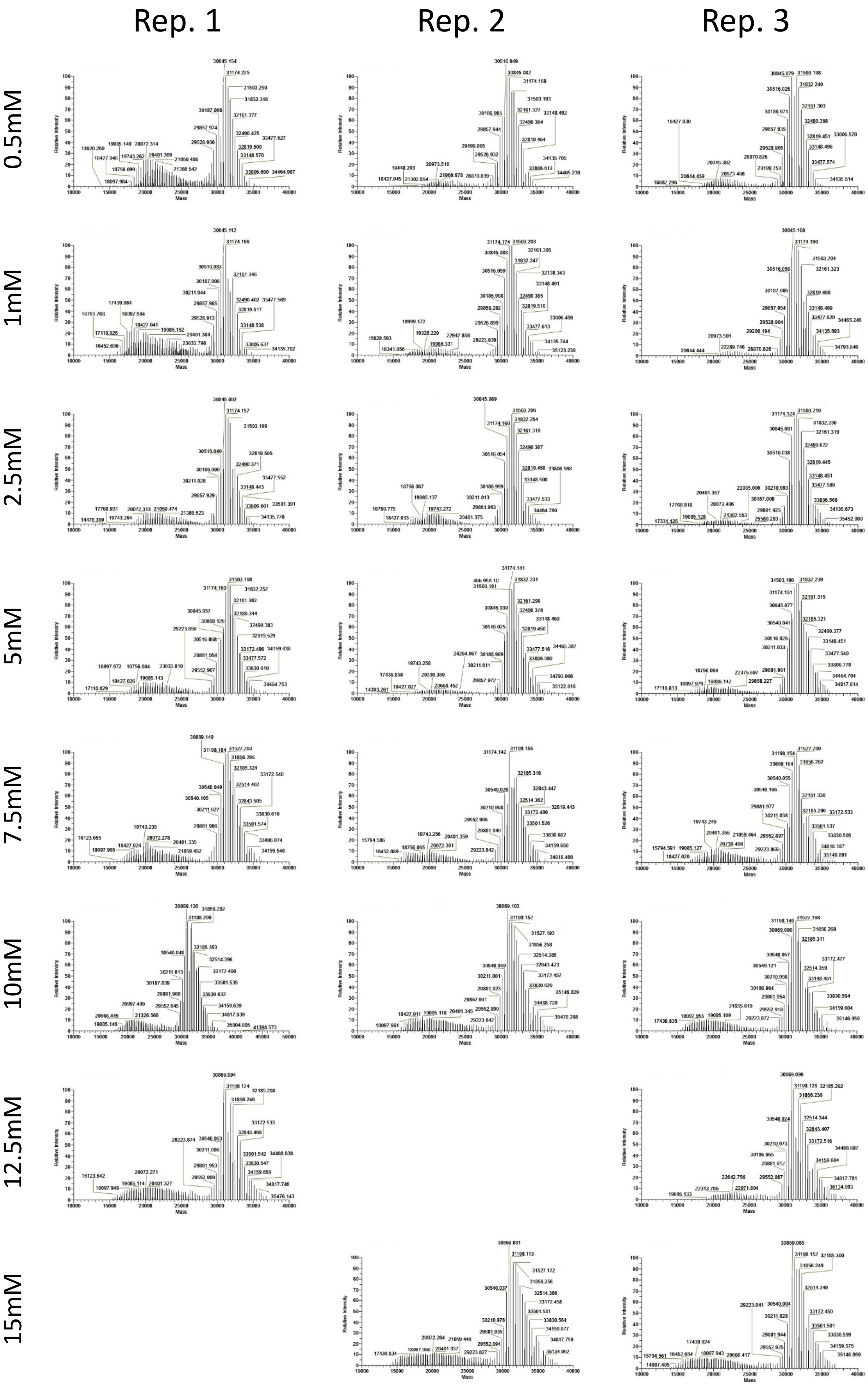


**Supplementary Figure S23. Template 6 mRNA ATP Concentration LC-MS Analysis.** Deconvoluted spectra of mRNA poly(A) tail species identified by LC-MS from RNase T1 digestion of *in vitro* transcribed material produced in IVT reaction with 0.5 mM, 1 mM, 2.5 mM, 5 mM, 7.5 mM, 10 mM, 12.5 mM or 15 mM ATP.

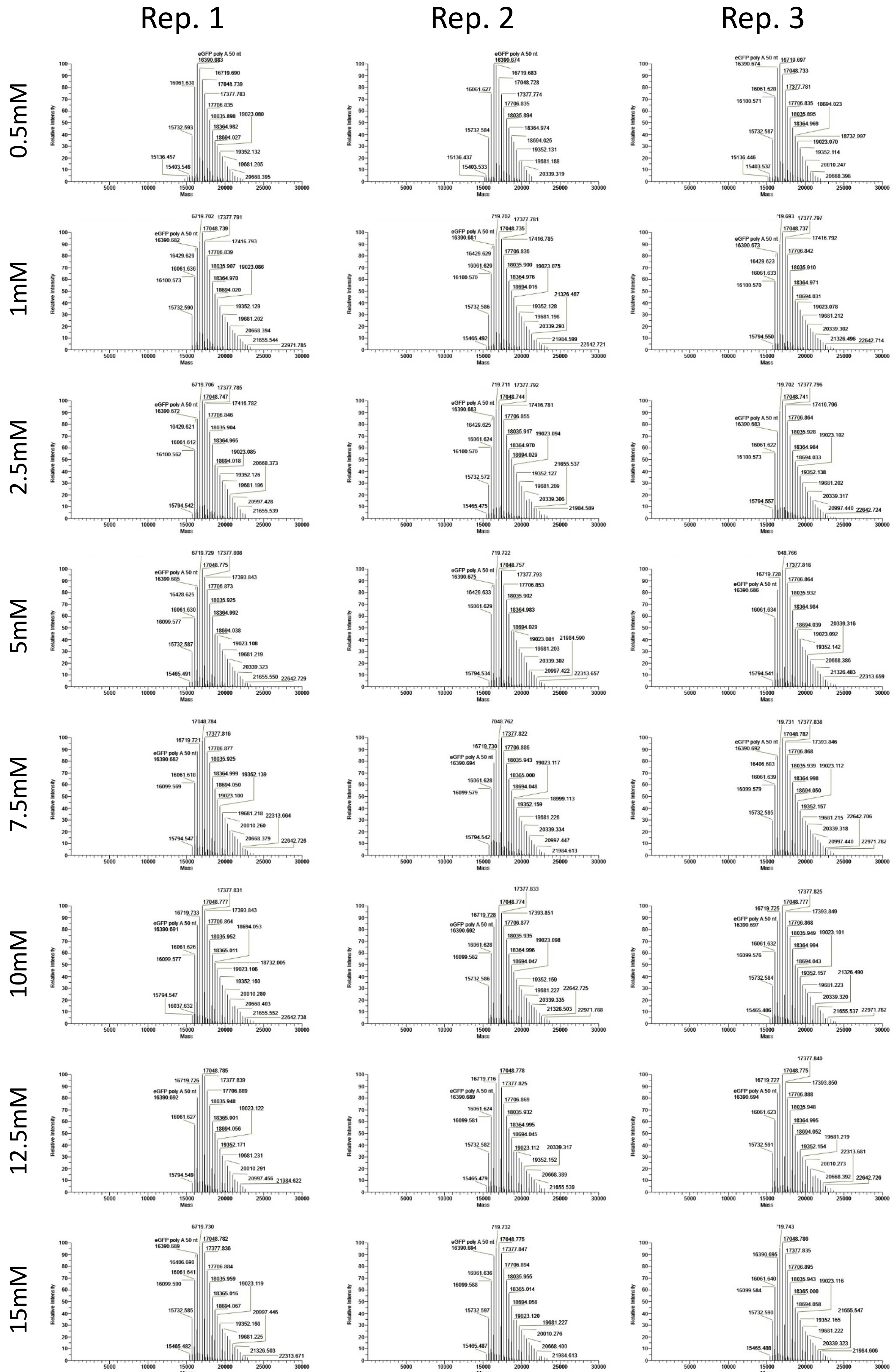


**Supplementary Figure S24. Template 2 mRNA CTP Concentration LC-MS Analysis.** Deconvoluted spectra of mRNA poly(A) tail species identified by LC-MS from RNase T1 digestion of *in vitro* transcribed material produced in IVT reaction with 0.5 mM, 1 mM, 2.5 mM, 5 mM, 7.5 mM, 10 mM, 12.5 mM or 15 mM CTP.

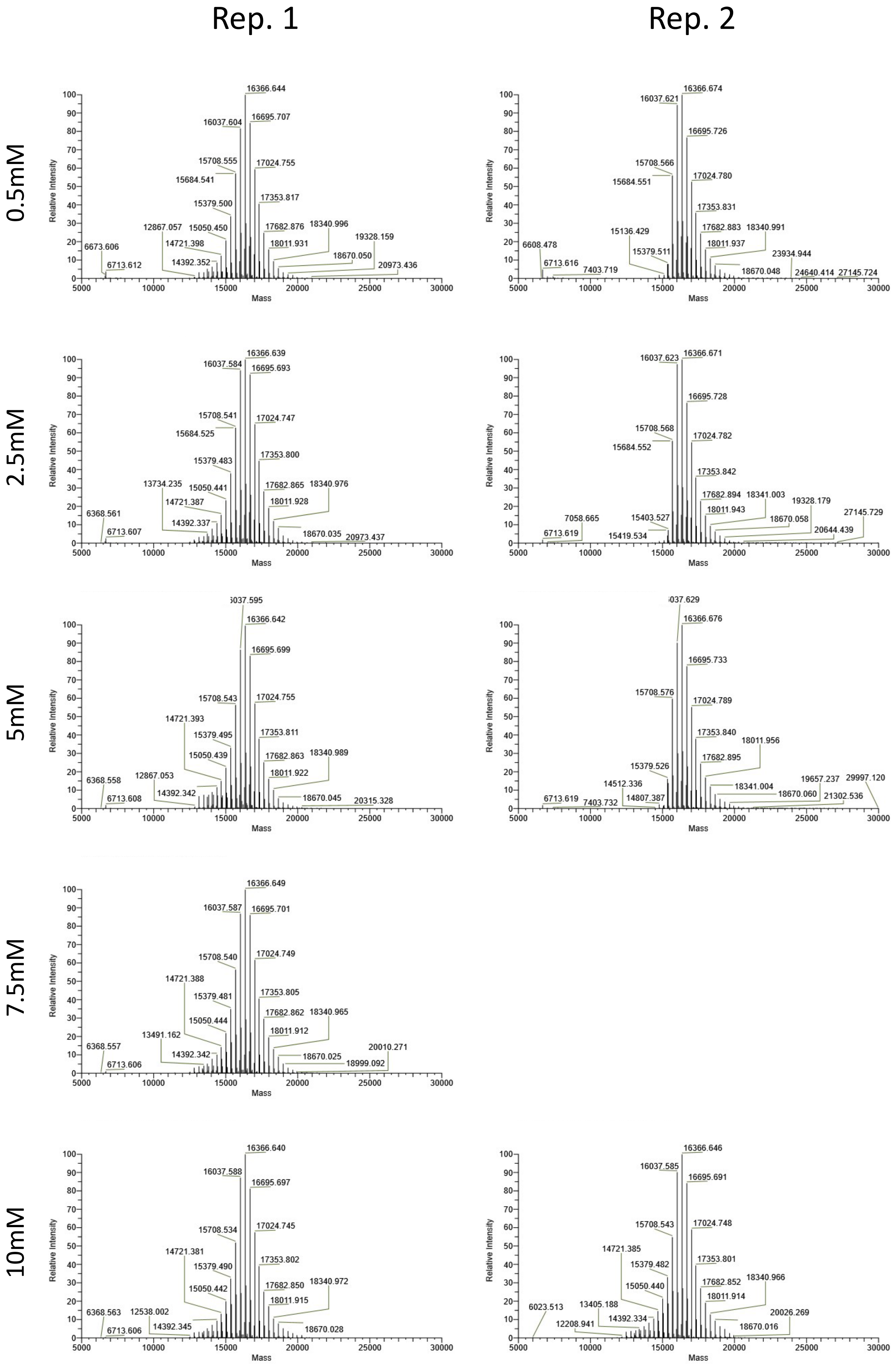


**Supplementary Figure S25. Template 2 mRNA UTP Concentration LC-MS Analysis.** Deconvoluted spectra of mRNA poly(A) tail species identified by LC-MS from RNase T1 digestion of *in vitro* transcribed material produced in IVT reaction with 0.5 mM, 2.5 mM, 5 mM, 7.5 mM, or 10 mM UTP.
